# A prevalent huge phage clade in human and animal gut microbiomes

**DOI:** 10.1101/2025.08.10.669567

**Authors:** LinXing Chen, Antonio Pedro Camargo, Yiting Qin, Eugene V. Koonin, Haoyu Wang, Yuanqiang Zou, Yi Duan, Hao Li

## Abstract

Huge phages are widespread in the biosphere, yet their prevalence and ecology in the human gut remain poorly characterized. Here, we report Jug (Jumbo gut) phages with genomes of 360-402 kilobase pairs that comprise ∼1.1% of the reads in human gut metagenomes, and are predicted to infect *Bacteroides* and/or *Phocaeicola*. Although three of the four major groups of Jug phages shared >90% genome-wide sequence identity, their large terminase subunits exhibited only 38– 57% identity, suggesting horizontal acquisition from other phages. Over 1,500 genomes of Jug phages were recovered from human and animal gut metagenomes, revealing their broad distribution, with largely shared gene content suggestive of frequent cross-animal-host transmission. Jug phages displayed high gene transcription activities, including the gene for a calcium-translocating P-type ATPase not detected previously in phages. These findings broaden our understanding of huge phages and highlight Jug phages as potential major players in gut microbiome ecology.

## Introduction

Viruses, in particular, those infecting bacteria and known as bacteriophages (phages), are the most abundant biological entities on Earth ^1^. The human gut contains a large diversity of phages ^2–4^, some of which have been reported to be widespread in the human gut ^5–7^, for example, crAssphage ^8,9^. Phages of the *Crassvirales* order (including crAssphage) that infect *Bacteroides* species ^10^ appear to be the most prevalent in the human gut, while also present in environmental viromes as well ^11^. These phages exhibit high phylogenetic diversity ^12,13^, and many members of the order possess unusual features such as interruption of genes by multiple self-splicing introns and inteins, and alternative genetic codes ^14^.

The composition of the human gut phageome has been shown to be unique and stable over the lifetime of individuals ^15^, but may shift during the early stage of life, although some phages can persist for a long time since the early-life periods ^16–19^. Composition shifts in the human gut phageome have been reported to be associated with chronic diseases ^20^, such as inflammatory bowel disease ^21,22^, malnutrition ^23^, and metabolic syndrome ^24^, suggesting that gut phages are relevant for human health.

The double-strand (ds) DNA phages of the *Caudoviricetes* class show a broad distribution of genome size, with the peak at about 50 kbp ^25^, whereas some, known as jumbo or huge phages, have genome sizes ≥200 kbp ^25–27^ and up to 841 kbp ^28^. Huge phages are diverse and widespread ^25^, and their large genomes with numerous genes encoding functionally diverse proteins provide a unique opportunity to study complex biological mechanisms in viruses. Examples of such mechanisms include genes encoding multiple components of the transcription and translation machineries^25^, enzymes augmenting various metabolic pathways of host cells, CRISPR-Cas systems along with other defense and counter-defense systems and more ^29,30^. Notably, some huge phages replicate their genomes within a nucleus-like structures that protect phage DNA against host defenses ^31–34^. The Lak phages, which infect *Prevotella* species, are among the very few huge phage clades that have been investigated in the human gut ^35,36^. They have a genome size of 540-660 kbp, which is comparable to that of some prokaryotes ^37^, and were detected mostly in gut metagenomes of humans on non-Western diets as well as various animals ^35,36^. In general, however, the prevalence, diversity, and ecological roles of huge phages in the human gut remain largely unknown.

In this work, we constructed the Huge Phage Genome Collection (HPGC) within which we identified a dominant huge phage clade, which we denoted Jug (after Jumbo gut) phages. The Jug phages were classified into four groups, and comparative genome analysis suggested distinct evolutionary scenarios. More than 1,500 Jug phage genomes were recovered from gut metagenomes of humans and various animals, demonstrating their broad diversity and distribution. Metatranscriptomics revealed the high activity of Jug phages in the human gut and provided information on cotranscription and RNA splicing of fragmented genes. We identified phage-encoded calcium-translocating P-type ATPases in Jug phages and other huge phages, an enzyme that has not been previously reported in phages, and showed that this gene was actively transcribed. The identification of Jug phages highlighted the capacity of HPGC for the discovery of ecologically important huge phage clades. Our analyses of the gene repertoires, distribution, transmission, and *in situ* activities of Jug phages document their potential importance as a major component of the human phageome and also expand our understanding of the ecology of huge phages in the gut microbiomes.

## Results

### The huge phage genome collection

We collected 10,690 viral genomes with a length of ≥200 kbp (Figure 1a, Supplementary Table 1), of which 9,372 were identified as huge phage genomes (see Methods), ultimately establishing a non-redundant dataset of 7,295 genomes (Supplementary Table 2). To our knowledge, this curated Huge Phage Genome Collection (HPGC) is the most comprehensive repository of huge phage genomes to date. The HPGC was dominated by genomes from the animal digestive system (2,288 genomes) and freshwater ecosystems (1,882 genomes). Additional sources included soil (936 genomes), marine systems (708 genomes), deep subsurface environments (345 genomes), and others (Figure 1b). About 70% of the genomes were complete or of high quality, with a strong size bias toward 200-400 kbp (Figure 1c).

**Figure 1.**
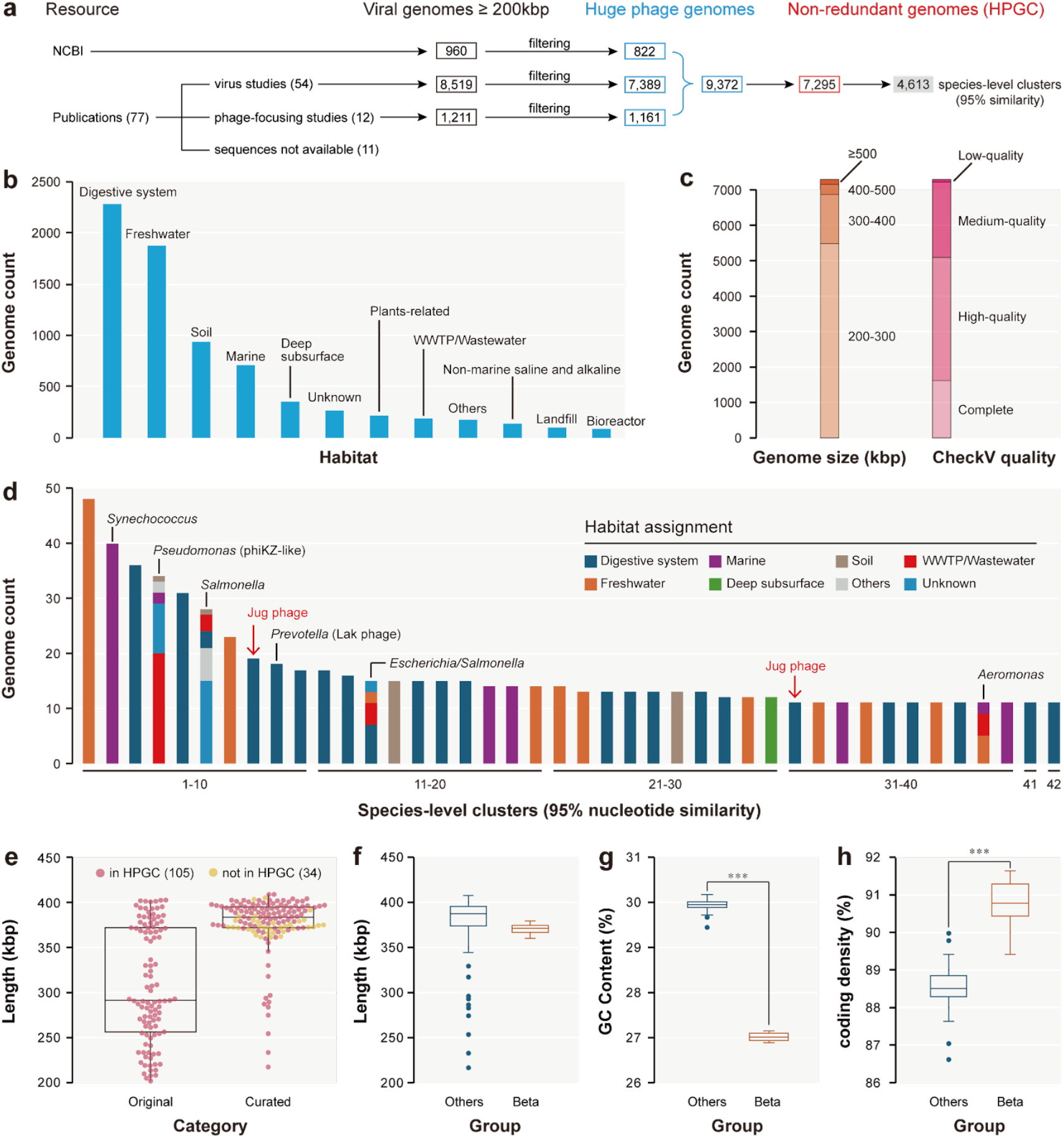
The construction of the Huge Phage Genome Collection (HPGC) and the identification of Jug phages. (a) The HPGC construction pipeline. (b) The habitat distribution. Only those habitats that contain 1% or more of the non-redundant genomes are shown. Otherwise, they are assigned to “Others”. Those without habitat information available were assigned to “Unknown”. (c) The length distribution and CheckV-based genome quality. (d) The large species-level clusters with >10 genomes. The bacterial hosts and the phage clade of the clusters are shown when available. The red arrows indicate the two species-level clusters of Jug phages. (e) The length of the Jug phage genomes. The original and curated lengths of the 105 HPGC genomes and the curated lengths of 34 newly assembled genomes are shown. (f)-(h) Comparison of the genomes from group Beta and the other three groups of Jug phages with respect to (f) genome length, (g) GC contents, and (h) coding density. The average values between group Beta and the others were compared using an unpaired t-test (*** P < 0.001).

Genome clustering at ≥95% nucleotide similarity identified 4,613 species-level clusters, with most clusters being singletons (3,547; Supplementary Table 2). Only ∼0.3% of the species-level clusters could be assigned to the taxonomic level of order based on geNomad analyses. Of the 42 large species-level clusters (>10 genomes each), 38 were detected in only one habitat each (Figure 1d). These large phage clusters included the phiKZ-like phages infecting *Pseudomonas*, the uncultured Lak phages infecting *Prevotella*, and phages infecting *Synechococcus, Salmonella, Escherichia*, and *Aeromonas*. Notably, 23 of the 42 large clusters included members from the animal digestive system, in particular, the human gut (Figure 1d). However, with the exception of the Lak phages, the diversity and potential ecological importance of these huge phages remained largely uncharacterized.

### A prevalent clade of huge phages from the human gut identified within HPGC

We focused on the human gut genomes within HPGC and clustered them at ≥90% nucleotide sequence identity (≥50% of genome length). The largest group contained 104 genomes (Supplementary Table 3), including the species-level clusters 08 and 31 (Figure 1d). We compared the amino acid sequences of the large terminase subunits (TerL) of these phages and divided them into two major groups, which we denoted Alpha and Delta. The TerL proteins from the two groups shared only ∼38% amino acid sequence identity. This observation motivated us to investigate the TerL identity gap. Searching the rest of HGPC identified one genome (huge_phage_7569), which shared ∼82% TerL identity to group Alpha, which we assigned to group Beta (Supplementary Figure 1). Subsequently, we performed a Pebblescout search ^38^ using the genomes of groups Alpha, Beta, and Delta, and identified a set of genomes that shared only ∼57% TerL identity but >90% nucleotide sequence identity with group Alpha (Supplementary Figure 2), which we named group Gamma. We referred to them as Jug phages after jumbo gut phages.

We manually curated 139 Jug phage genomes (Supplementary Table 3), including 96, 27, 4, and 12 genomes in groups Alpha, Beta, Gamma, and Delta, respectively. Manual curation improved the quality of the 105 genomes within HPGC by extending their length from 307 kbp to 375 kbp on average, with all 34 newly assembled genomes being >360 kbp (Figure 1e). Compared with the other three groups, group Beta Jug phages had smaller genome sizes (Figure 1f) and lower GC contents (27% vs 30% on average; Figure 1g), but higher coding density (90.7% vs 88.6% on average using code 11; Figure 1h). The Jug phage genomes encompassed from 320 to 555 (mean 501) predicted protein-coding genes. We annotated the genes using both Pfam domain search and protein structure prediction and comparison, and identified many of the viral marker genes, which indicated Jug phages are T4-like phages (Supplementary Table 4).

All the curated genomes of groups Alpha, Gamma, and Delta were from the human gut. In contrast, the Beta group had a broader animal-host range, with members with curated genomes identified in the guts of humans, dogs, cats, chickens, swans, and quails (Supplementary Table 3), and also in ducks with fragmented genomes (Supplementary Figure 3).

### Phylogeny, genome comparison, and the core gene set of Jug phages

Groups Alpha and Beta shared the highest TerL identity, whereas the comparison of genome nucleotide sequences, as well as amino acid sequences of DNA-directed RNA polymerase subunit beta’ (RpoC), major capsid protein (MCP), and DNA polymerase showed higher similarity between groups Alpha, Gamma, and Delta (Figure 2a). These observations were supported by genome clustering (Supplementary Figure 4) and the detection of a large number of single-copy genes shared by the three groups (406 in total, of which 371 had average pairwise identity >90%; Supplementary Table 5).

**Figure 2.**
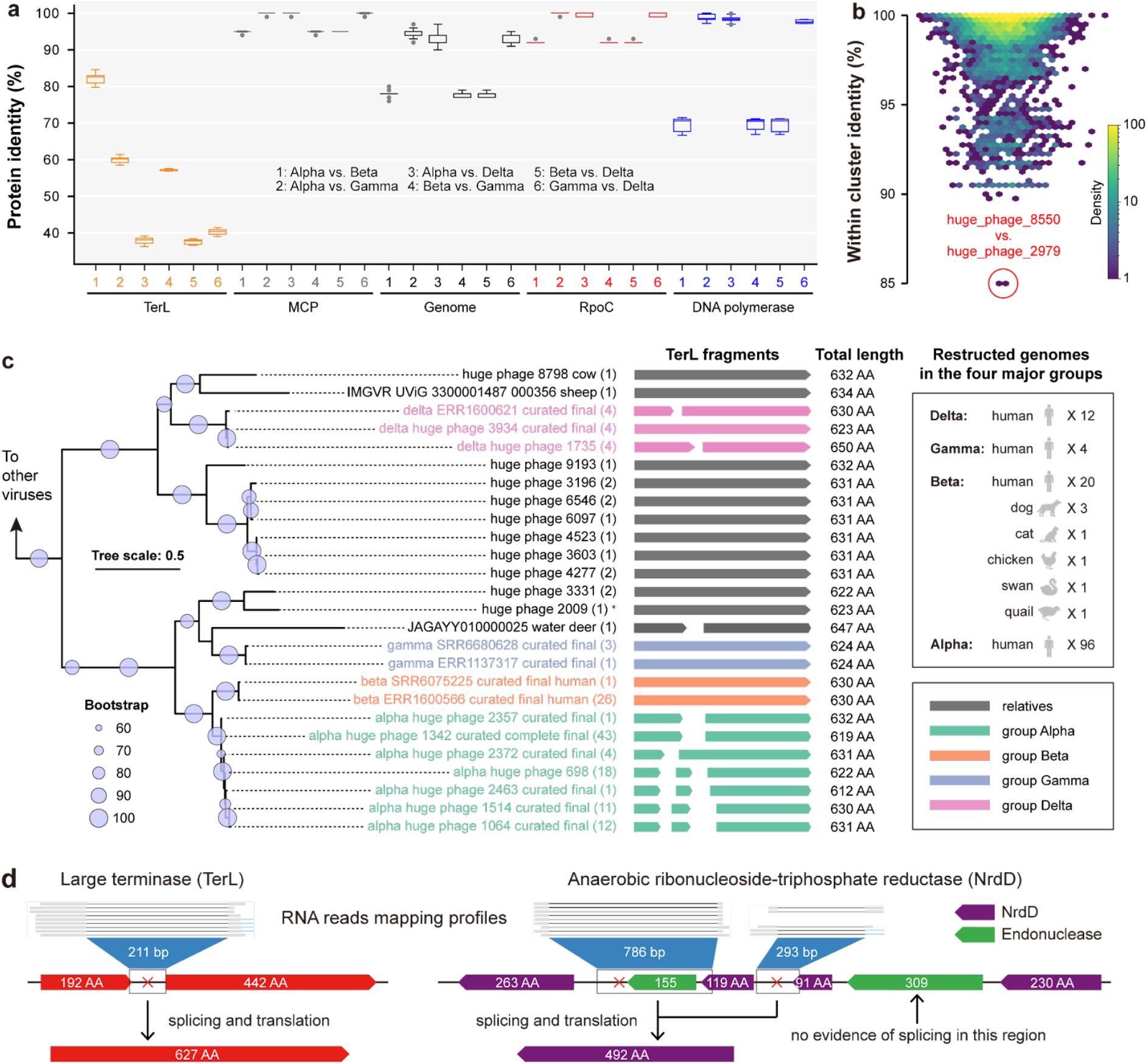
The phylogeny and genomic features of the Jug phages. (a) Identity of TerL, MCP, genome, RpoC, and DNA polymerase between groups. The outliers are indicated by gray circles. (b) TerL identity within other HPGC clusters (≥90% nucleotide similarity) with at least 5 members. The cluster with the lowest TerL identity is indicated with a red circle. (c) The TerL-based phylogeny. The TerL fragments of each curated Jug phage genome were concatenated individually and clustered at ≥99% identity. The TerL sequences of relatives were obtained from HPGC and IMG/VR via BLASTp and clustered at ≥99% identity as well. Only the cluster representatives were included, and the count of proteins in each cluster is indicated in the brackets. The total length of the TerL fragments is shown. All the corresponding genomes were from the gut microbiomes except one from a wastewater sample (indicated by *). (d) The RNA reads mapping evidence of cotranscription and RNA splicing of split genes. Examples of *terL* and *nrdD* genes are shown. Note that the *nrdD* gene was split into 4 fragments, and no mRNA splicing evidence was observed for the region between the 230AA and 91AA fragments.

Altogether, we detected 39 single-copy genes with ≥70% protein identity that are conserved in all four Jug phage groups (Supplementary Figure 5, Supplementary Table 6) and formed several conserved gene arrays (Supplementary Figure 6). These conserved genes included those for MCP, anaerobic ribonucleoside-triphosphate reductase activating enzyme (NrdG), DNA topoisomerase IV, translation initiation factor 1A, and Ribonuclease H, with most of the others without functional annotation. Notably, other viral marker genes, such as phage portal protein and prohead protease (Supplementary Table 4), were fragmented (see below) and/or highly divergent and thus not counted as single-copy core genes.

Given the low TerL identity contrasting the high (>90%) genome sequence identity among groups Alpha, Gamma, and Delta (Supplementary Figure 4), we searched the entire HPGC for other clusters of genomes with a similar pattern of sequence conservation. However, no other genome clusters in the HPGC dataset exhibited TerL identity <85% (Figure 2b, Supplementary Figure 7), suggesting that the “high genome but low TerL identity” pattern is unique to Jug phages. Phylogenetic analyses of TerL proteins showed that group Gamma was most closely related to a huge phage from the water deer gut, whereas group Delta clustered with huge phages from the guts of ruminants such as sheep and cows (Figure 2c, Supplementary Figures 8-10). This pattern suggests independent acquisition of *terL* genes from gut-resident huge phages outside the Jug group, which was supported by the GC content profiles of the *terL*-encoding regions (Supplementary Figure 11). Moreover, the genes immediately upstream of *terL* across the Alpha, Gamma, and Delta groups that typically encode predicted endonucleases were highly divergent, suggesting that these genes could have been transferred together with *terL* (Supplementary Figure 12).

The relatively lower genome similarity between groups Alpha and Beta challenged their phylogenetic relatedness despite the high identity of their TerL sequences, which could be due to HGT as well. We explored the rest of HPGC and did not detect any other genomes with >50% genome identity to group Beta. On the other hand, groups Alpha and Beta shared ∼95% MCP identity (Supplementary Table 7). The MCP phylogeny supported the close relatedness of groups Alpha and Beta (Supplementary Figure 13).

### Jug phage genes are frequently interrupted by group I introns and putative inteins

During the curation of the Jug phage genomes, we noticed that many genes appeared to be fragmented. A search for self-splicing introns (see **Methods**) led to the identification of 2,513 group I introns in 136 of the 139 curated Jug phage genomes (18.5 introns, on average, in each genome; Supplementary Table 8), with fewer introns in group Beta compared to the other groups (Supplementary Table 9). These introns were on average ∼206 nt in length, suggesting that they were the small internal group I-like ribozymes ^39^. In addition, we identified numerous homing endonucleases of different families, in particular, 505 HNH, 80 PDDEXK_5, 55 GIY-YIG, and 17 LAGLIDADG endonucleases, as well as 26 Intein_splicing domain-containing proteins (Supplementary Table 4). Among them, except for 88 HNH and 9 LAGLIDADG domains that were inside other genes, all the remaining were stand-alone genes.

The group I introns and genes encoding various homing endonucleases interrupt many essential genes of the Jug phages. For example, the *terL* genes in all Alpha and most Delta genomes were split into 2-3 fragments due to group I intron insertion (Supplementary Figure 14). The *nrdD* genes encoding anaerobic ribonucleotide reductase were fragmented in 95/104 genomes; all the intact *nrdD* genes were exclusively from group Beta (Supplementary Table 10). Insertions between the fragments commonly encoded endonucleases or proteins with “Intein_splicing” domains. In addition, DNA polymerase genes were also frequently split by intervening endonuclease sequences (Supplementary Table 11).

To validate the predicted group I introns in Jug phages, we re-analyzed three gut samples of three male adults containing Jug phages for which metatranscriptomes were available ^40^, given that there was no metatranscriptome for any of the 139 curated Jug phage genomes (see **Methods**). Analysis of these three Jug phage genomes and the corresponding metatranscriptomes showed that for 37 of the 76 predicted group I introns, splicing was supported by metatranscriptome reads mapping (Supplementary Table 12). For example, the split *terL* and *nrdD* genes were cotranscribed as an operon (Supplementary Figure 15). Some of the RNA reads were precisely mapped to the flanking regions of the introns (Figure 2d), directly confirming the restoration of the respective coding regions by splicing (Supplementary Figure 16).

### Jug phages infect *Bacteroides* and/or *Phocaeicola* species

The iPhoP analysis indicated that Jug phages infect *Bacteroides* and/or *Phocaeicola* (Supplementary Table 13), which was supported by local CRISPR-Cas spacer analysis, with a total of 34 unique targeting spacers (Supplementary Figure 17, Supplementary Table 14). *Bacteroides* and *Phocaeicola*, along with *Prevotella*, belong to the Bacteroidaceae family (GTDB taxonomy) and are typically abundant in the human gut. Previously, the *Bacteroides*-infecting crAss-like phages (89-192 kbp in length) ^8,13^, and the *Prevotella*-infecting Lak phages (>540 kbp in length) ^35,36^ have been reported to be widespread in the human gut. However, Jug phages showed no significant similarity to any of these phages apart from sharing several hallmark genes (see **Methods**).

To further validate the predicted host-virus relationship, we performed co-occurrence analyses for the 125 metagenomic assemblies that were used for the manual curation of Jug phage genomes. In these metagenomes, we identified 28 *Bacteroides* and 16 *Phocaeicola* species, with at least one of these represented in each of the 125 samples. However, only *P. vulgatus* and *B. clarus* were detected in more than half of the samples (Figure 3a). In one sample (SRR7403886), we detected one *Bacteroides* species without *Phocaeicola*, suggesting its potential host-phage relationship with the co-occurring group Gamma Jug phage. We attempted to confirm specific host-virus relationships by investigating the samples detected with only 2 or 3 *Bacteroides*/*Phocaeicola* species, but failed to construct any for them (Supplementary Figure 18).

**Figure 3.**
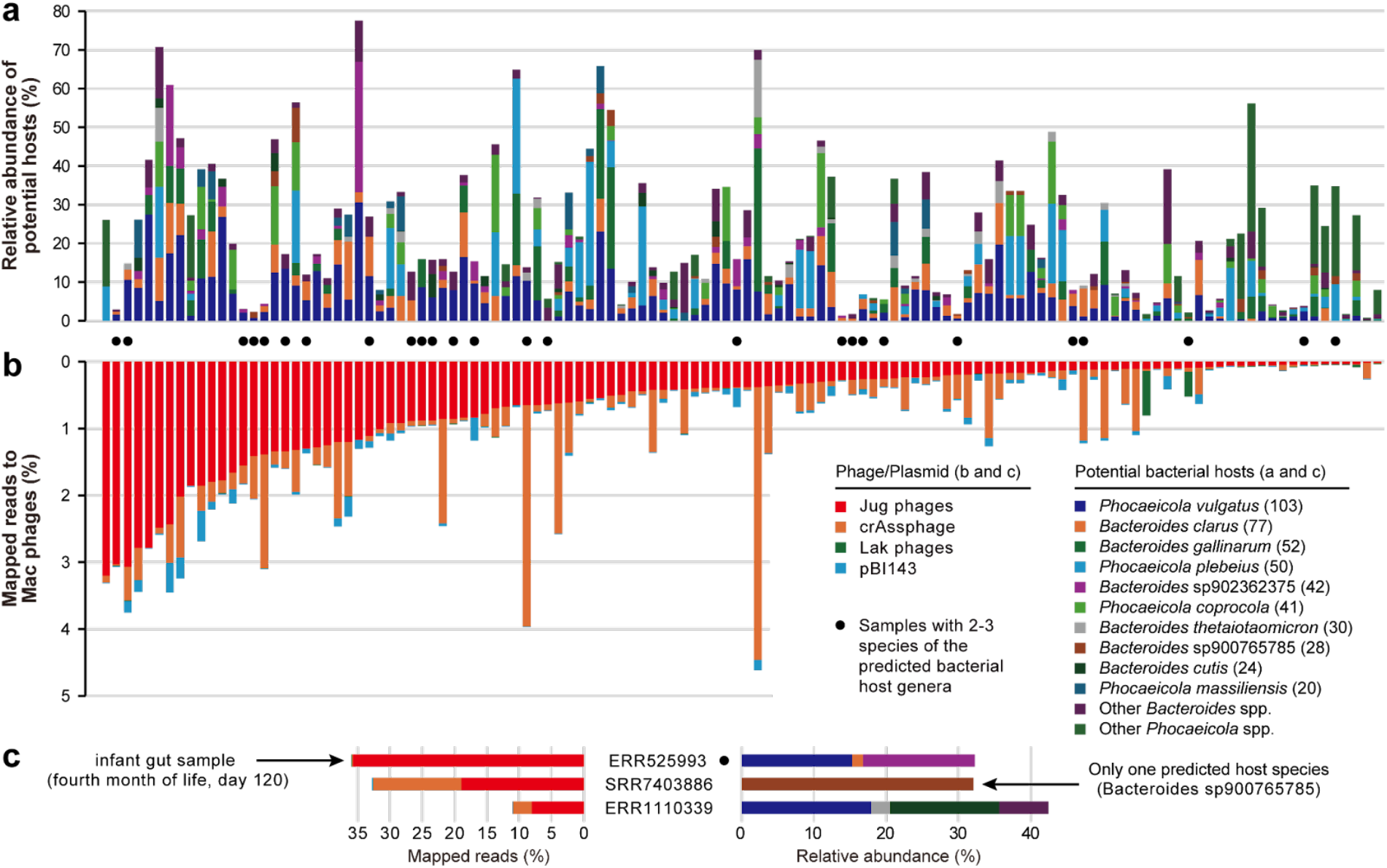
The predicted bacterial hosts of Jug phages and host-phage co-occurrence. (a) The relative abundance of predicted hosts in the samples. The numbers in the brackets following the species names indicate the count of samples with detection. Only those species detected in ≥20 samples are listed, and the rest are assigned to “other species”. (b) Percentage of mapped metagenomic reads. pBI143 is a widespread plasmid in the human gut ^44^. The samples with 2 or 3 predicted host species detected are highlighted with black circles, with further analyses shown in Supplementary Figure 18. (c) The three samples with >8% of metagenomic reads mapped to the Jug phage genomes. The data for the sample with the highest fraction of mapped reads and the sample with only one predicted bacterial host species are shown. One of the three samples contained only one *Bacteroides* species without *Phocaeicola*; thus, the specific host-virus relationship was tentatively inferred.

On average, Jug phages accounted for 1.1% of the metagenomic reads, with >8% in three bulk metagenomic samples (Figures 3b and c). One sample, from an infant in the fourth month of life, contained a group Alpha Jug phage, which accounted for ∼35.8% of the mapped reads. This particular gut microbiome contained 14 bacterial species, including *Bacteroides clarus, Bacteroides* sp902362375, and *Phocaeicola vulgatus*, with a cumulative relative abundance of 29.7% for these three species (Figure 3c, Supplementary Table 15). We identified the same Jug phage in the gut microbiomes of the mother at delivery, and the infant at birth and in the 12th month (Supplementary Figure 19), suggesting transmission of the Jug phage from the mother to the infant.

### Jug phages are widespread in the guts of humans and animals

To assess the diversity and distribution of Jug phages, we searched the Logan assembly dataset ^41^ using *terL* as a marker and compared the retrieved contigs against the curated Jug phage genomes (Figure 4a). We identified 2,733 samples containing Jug phage-related contigs ≥1 kbp and a cumulative phage genome length ≥200 kbp (see **Methods**), with nearly all derived from gut microbiomes except for 12 from wastewater (Supplementary Table 16). The wastewater-derived genomes were highly fragmented (Supplementary Figure 20).

**Figure 4.**
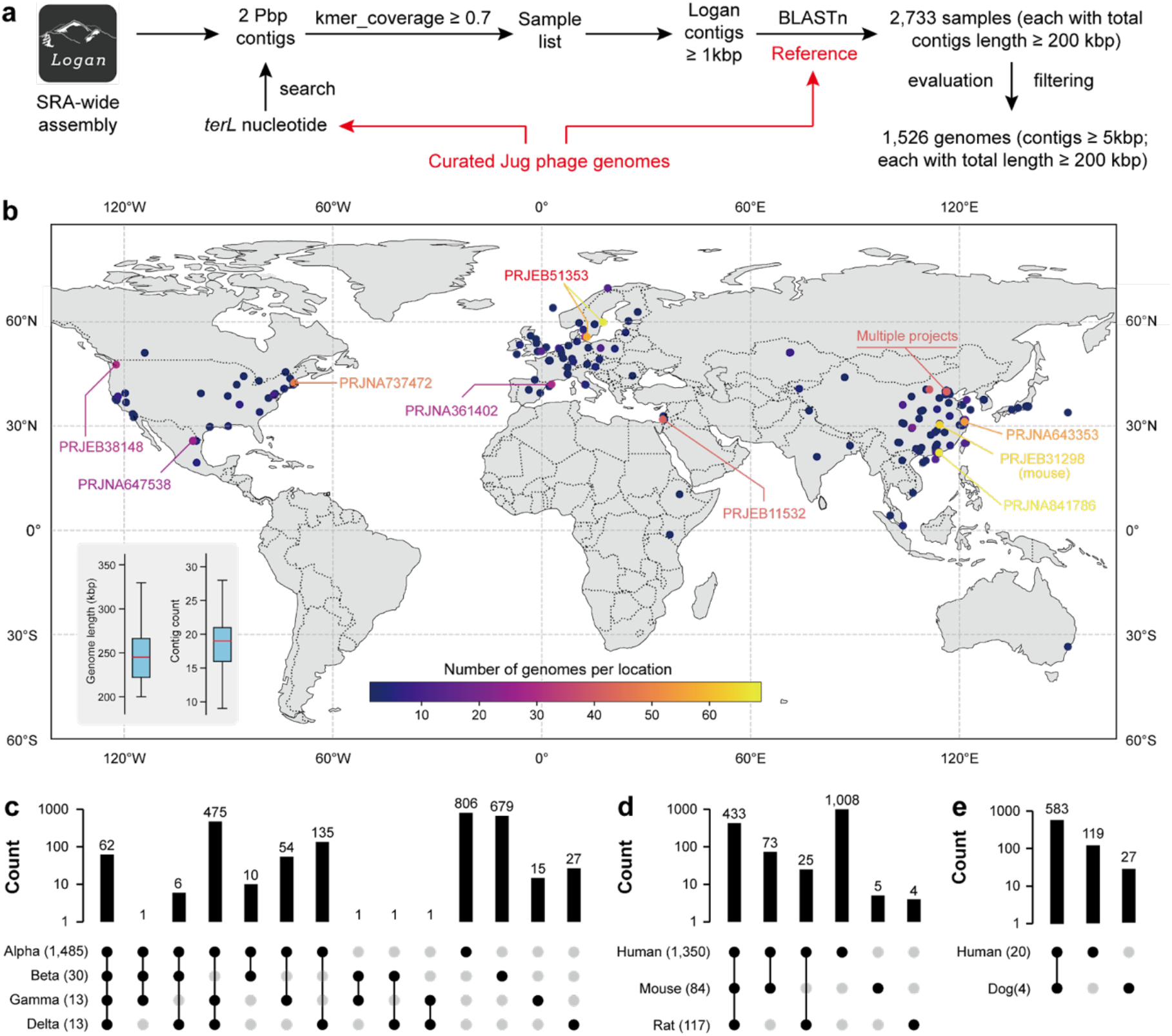
Cluster analysis of Jug phage proteins. (a) The Logan-based analysis pipeline to obtain Jug phage genomes. Each of the BLASTn hits was filtered to allow ≥90% nucleotide identity across ≥90% of the contig length when matched to a curated Jug phage genome. (b) The global distribution of Logan-retrieved Jug phage genomes. 1,422 out of the 1,523 genomes (Supplementary Table 17) have location information and are shown on the map. All the genomes have a minimum length of 200 kbp, with all contigs in the genomes ≥5 kbp in length. The locations with more than 30 genomes retrieved were highlighted with the corresponding NCBI project IDs shown (all projects but one were from the human gut). The insertion in the lower-left corner shows the total length and contig count in each retrieved genome. Some regions of the world map are not shown as there was no genome identified or due to low map resolution. The distribution of protein clusters within (c) the four Jug phage groups, (d) human, mouse, and rat of group Alpha, and (e) human and dog of group Beta. In (c), (d), and (e), the shared or specific protein clusters are indicated by bars, with the count of the protein clusters shown. The numbers of Jug phage genomes in each group or animal host are shown in the brackets. The protein clusters are defined using CD-HIT at ≥70% protein identity (see **Methods**).

Predicted proteins from the Logan contigs were compared to the proteins of curated Jug phage genomes (TerL, MCP, RpoC, and DNA polymerase), and a single copy of each gene was detected in nearly all samples except one (NCBI SRA = SRR10680429), which contained one Alpha and one Beta member. These findings indicated that gut microbiomes typically harbor only one dominant Jug phage genotype. We thus reconstructed draft genomes from the individual samples by binning the matched contigs ≥5 kbp, yielding 1,526 genomes ≥200 kbp in length (Figure 4a), the majority (1,450) belonging to group Alpha. These draft genomes comprised 9–29 contigs (Figure 4b, Supplementary Table 17) and showed 99.5% average nucleotide identity to the curated Jug phage genomes (Supplementary Table 18). Notably, group Alpha Jug phages were also found in non-human animal hosts, including mice (84 genomes), rats (51 genomes), and hamsters (3 genomes).

We identified 2,273 protein clusters in the curated and Logan-retrieved Jug phage genomes. Only ∼10.9% of the cluster representatives were assigned known functions, with Pfam domains such as NRDD, AAA, Phage_lysozyme2, Amidase_3, CLP_protease, DNA_topoisoIV, eIF-1a, and HNH_3 among the most frequently detected (Supplementary Table 19). Group Alpha shared numerous protein clusters with groups Gamma and Delta, but few with group Beta (Figure 4c). Over 35% of the protein clusters were unique to group Alpha. Within this group, human-, mouse-, and rat-derived Jug phages shared 433 protein clusters, with only five and four clusters unique to mouse and rat, respectively (Figure 4d). Similarly, within group Beta, Jug phages from the dog gut metagenomes shared >95% of proteins with those from humans (Figure 4e). These shared clusters generally exhibited high sequence identity (Supplementary Figure 21), and phylogenetic analyses showed no animal-host-specific clades (Supplementary Figure 22). These findings suggest frequent transmission of Jug phages between animal hosts.

### Jug phages are transmissible among humans and sensitive to dietary intervention

To explore the transmission of Jug phages, we re-analyzed the human gut samples of donors and recipients of a study of fecal microbiota transplantation (FMT) for the cure of ulcerative colitis ^45^. We confirmed that the recipients harbored no Jug phages before FMT (Supplementary Figure 23, Supplementary Table 20). Following FMT, Jug phages identical to those in the donors were identified in the guts of 7 recipients during the treatment and were retained for up to 14-25 weeks. These results suggested that the Jug phage could be transmitted among humans.

To investigate the potential effects of diet on Jug phages, we re-analyzed the human gut samples of overweight or obese participants in two dietary intervention studies (study 1 and study 2) ^46^. The participants were given fibre-containing snacks while consuming meals that are high in saturated fats and low in fruits and vegetables (HiSF-LoFV) (Supplementary Figure 24, Supplementary Table 21). Group Alpha Jug phages were identified in participant S11 samples of both studies 1 and 2, and in participants S13 and S15 samples of study 1. The Jug phages were present in all the 58 analyzed samples, albeit with very low coverage in 8 samples (Supplementary Figure 25). *P. vulgatus* and *Bacteroides* sp902362375 were the only two shared potential host species identified in all three participants. We did not observe any significant correlation between the abundances of these bacteria and the Jug phages. However, the abundance of the Jug phages increased towards the end of the continuous consumption of fiber snacks as supplements in all three participants, suggesting that fibre-enriched diets promoted the reproduction of the Jug phages, conceivably, by enhancing the growth of the host bacteria.

### Jug phages are transcriptionally active in the human gut

To investigate the *in situ* activities of Jug phages, we re-analyzed metagenomic and metatranscriptomic datasets from three adult male gut samples that harbored Jug phages (see above) ^40^. All three Jug phages (one per individual) were from group Alpha (Supplementary Table 22) and accounted for 3-59% (26% on average) of all the virus-retrieved DNA reads (Figure 5a). Notably, Jug phages were highly active in 7 of the 12 analyzed samples, with their transcripts accounting for 56-96% (79% on average) of all virus-retrieved RNA reads, which was by far the highest among all viruses co-occurring in these samples. In these 7 samples, 87-99% of the Jug phage genes were expressed (Figure 5b). Genes for proteins involved in lytic infection, in particular, virion formation, such as TerL, MCP, portal protein, prohead protease, baseplate proteins, tail components, and tape-measure proteins, ranked relatively low in transcriptional activity (Supplementary Table 23), suggesting that the majority of the Jug phages were not involved in active lysis of the host cells at the time of sampling.

**Figure 5.**
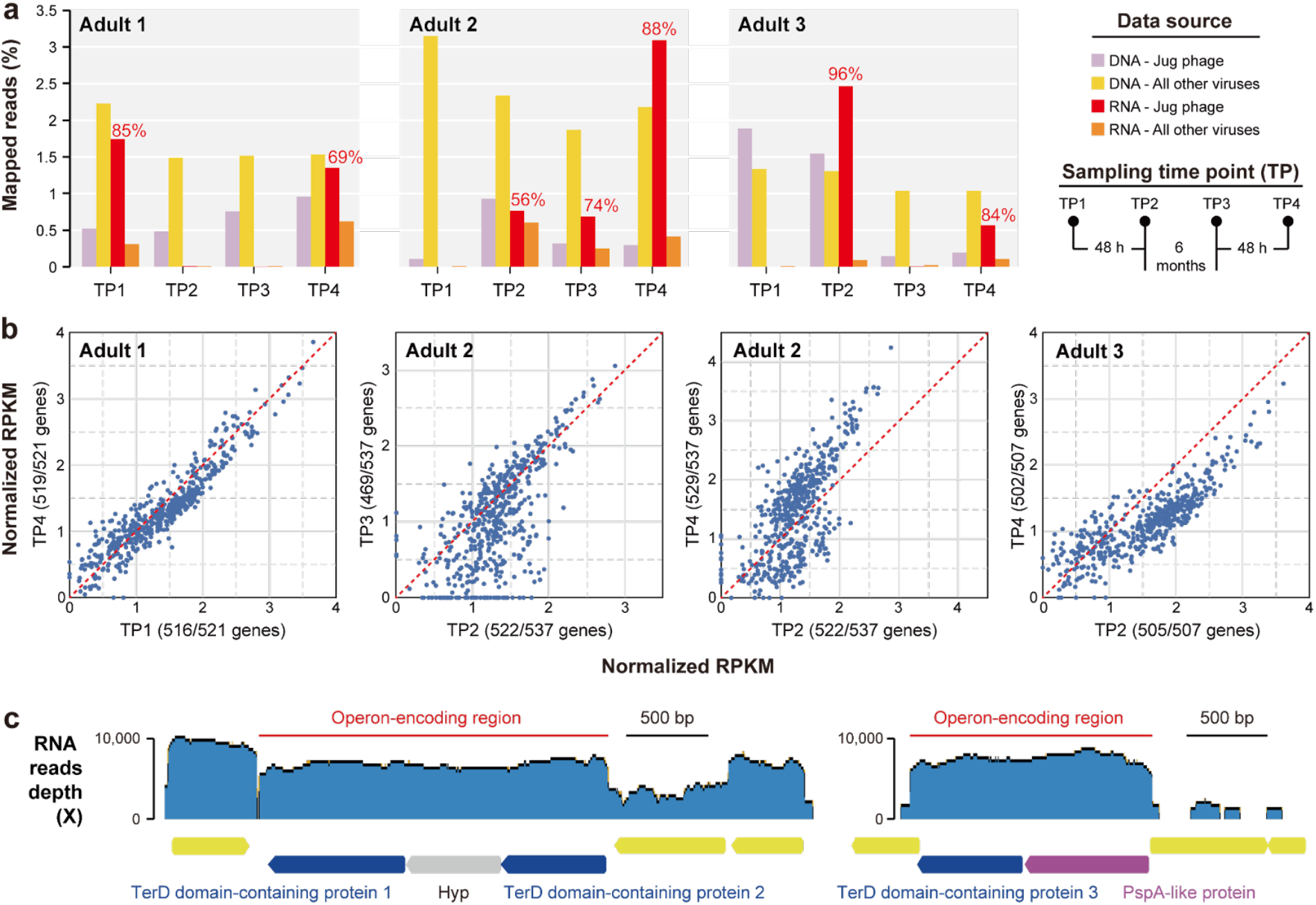
Transcription activity of Jug phages in the guts of three adult men. (a) The relative abundance of DNA and RNA reads mapped to the Jug phages and all other viruses in the corresponding samples. For the 7 samples with transcriptionally active Jug phages, the percentages of RNA reads mapped to the Jug phages against the total RNA reads mapped to all viruses are shown above the bars. The sampling procedures are shown to the right in brief. (b) The scatter plots compare the transcriptional levels of protein-coding genes in the Jug phages of the three adults. The number of transcribed genes in each time point sample is indicated in the bracket (see Supplementary Table 23 for details). Each blue circle represents a gene; the closer the circle to the red dashed line, the more similar the transcriptional activity is in the two samples. (c) The transcriptional profiles of the genes for TerD domain-containing proteins and others. The Jug phage genome from adult 1 is shown as an example. The RNA reads were mapped to the genome. The operon-encoding regions (suggested by the RNA reads mapping profiles) are indicated by the red lines. Hyp, hypothetical protein.

Each of the three Jug phage genomes contained three *terD* genes: two flanking a hypothetical gene without predicted function, and a third adjacent to a *pspA*-like gene (Figure 5c). Transcriptional profiling showed that each region was independently transcribed as an operon. Given that TerD-related proteins typically participate in stress response ^47^, the high RNA read mapping depth of the TerD-encoding regions suggests that Jug phages might experience strong environmental stress at the time of sampling. Among the Jug phage protein clusters, 9 consisted of TerD-related proteins (>2,200 proteins, ∼0.45% of all predicted proteins; Figure 4c). We found that TerD-related proteins are widely encoded by gut bacteria, and those from the same genus generally clustered together in the phylogenetic tree (Supplementary Figure 26). Furthermore, TerD-like proteins encoded by Jug phages grouped closely with those from *Bacteroides, Phocaeicola*, and *Prevotella*, suggesting the possibility of HGT of these stress-related genes between Jug phages and their hosts.

### A calcium-translocating P-type ATPase identified in Jug and other phages

Notably, among the other highly transcribed genes was a gene encoding a calcium-translocating P-type ATPase (Supplementary Table 23). This gene was identified in the majority of the curated (60.4%; all four groups) and the Logan-retrieved (74.1%) Jug phage genomes. Protein structure prediction and comparison confirmed the annotation (Figures 6a-c, Supplementary Figure 27), and sequence alignment showed the 10 transmembrane helices typical of P-type ATPases and the key functional residues (Supplementary Figure 28). Many of the gut microbes encode this ATPase, which is an integral membrane protein that couples ATP hydrolysis to the export of Ca^2+^ ions, thereby maintaining low cytoplasmic Ca^2+^ levels essential for calcium homeostasis ^48^.

**Figure 6.**
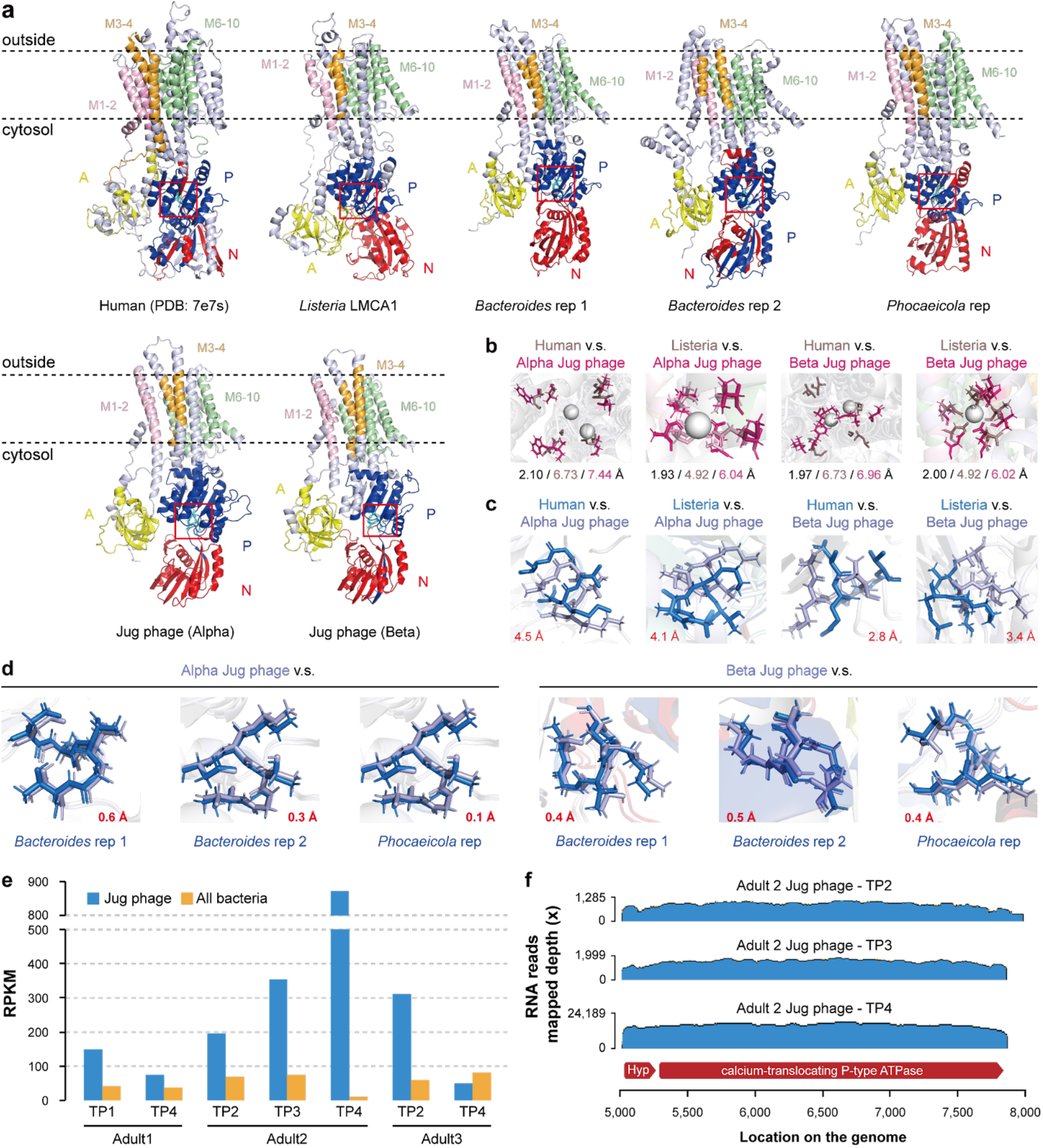
A calcium-translocating P-type ATPase identified in phages. (a) The protein structure of the human calcium-translocating P-type ATPase (PDB: 7e7s), and the predicted models of the ATPases from *Listeria monocytogenes* (LMCA1), *Bacteroides* and *Phocaeicola* species, and Jug phages. In the structure representations, “A” stands for the actuator domain, “P” stands for the phosphorylation domain, and “N” denotes the nucleotide-binding domain. The dashed lines indicate the membrane. The transmembrane helices are highlighted with different colors. The conserved phosphorylation motif “DKTGT” is highlighted in cyan and displayed as sticks. (b) Superposition of the calcium coordination sites. The sum distance between the Ca^2+^ (gray spheres)-binding residues between the ATPases from Jug phage and human (or bacteria) (shown in black), the sum distance of the Ca^2+^-binding residues to the Ca^2+^ ion in the ATPases from human (or bacteria) (shown in gray), and the sum distance from Ca^2+^-binding residues to the Ca^2+^ ion in the ATPase of Jug phage (shown in pink) are shown at the bottom. Note that the human ATPase can coordinate two Ca^2+^ ions. (c) Superposition of the “DKTGT” motifs from the Jug phage and human and bacterial P-type ATPases. The location of the motif in the protein is indicated by red boxes in (a). The distance (Å) between the sapratet (D) residue from the Jug phage and the reference ATPase is shown in red. (d) Superposition of the “DKTGT” motif between the ATPases of the Jug phages and predicted bacterial hosts. The distance between the D residues is shown in red. (e) The normalized transcriptional activities (RPKM) of the calcium-translocating P-type ATPase genes in Jug phages and the co-occurring bacteria in the gut microbiomes. The RPKM values of all bacteria were aggregated. (f) Co-transcription of the calcium-translocating P-type ATPase gene and the adjacent upstream gene.

Phylogenetic analysis showed that the P-type ATPases of Jug phages were most closely related to those from *Bacteroides* and *Phocaeicola* species (Supplementary Figure 29), suggestive of HGT and supporting the bacterial host prediction for Jug phages. The arrangement of the aspartic acid residues (“DKTGT”), which are phosphorylated during the catalytic cycle, was highly consistent between the P-type ATPases of Jug phages and their predicted hosts as well (Figure 6d). Surprisingly, in all but one of the 7 samples, the transcriptional activity of the Jug phage-encoded ATPase gene was higher than the cumulative activities of those encoded by all the co-existed bacteria in the gut microbiomes (Figure 6e).

To date, the only viral calcium-translocating P-type ATPase was documented in *Nucleocytoviricota* viruses infecting *Chlorella*, where this gene has been shown to be actively transcribed during infection ^49^. Our extended analysis identified this ATPase in 81 other HPGC huge phages (200-447 kbp in length) and 4 smaller-sized phages (159-197 kbp in length) (Supplementary Table 24). Notably, all these phage genomes were reconstructed from gut microbiomes, and the phage-encoded ATPases were most similar to those from Bacteroidaceae (including *Bacteroides* and *Phocaeicola*) or Bacillota (Supplementary Figure 30).

We found that the gene immediately upstream of the calcium-translocating P-type ATPase was co-transcribed with the ATPase gene (Figure 6f). This adjacent gene is present in all Jug phages encoding the ATPase, and encodes a small protein (77-91 amino acids) without detectable homologs (Supplementary Figure 31). The consistent presence of the two genes as an operon in the Jug phages implies that the small protein is functionally linked to the ATPase, perhaps regulating its activity.

## Discussion

Given their pivotal roles in shaping microbial community composition, in particular, human-associated microbiomes, element cycling, and the evolution of their hosts, phages have lately attracted renewed, strong interest in microbiology research ^50^. Huge phages are particularly notable because of their large genomes that encompass genes for complex biological mechanisms that are usually absent in smaller-sized phages ^25,51^. Our understanding of huge phages has dramatically extended in the past several years, especially with the development of genome-resolved metagenomics and related analysis tools ^52–61^. However, the absence of a curated reference dataset of huge phages impedes deeper ecological insights. To fill this gap, we constructed HPGC. While conventional phage studies routinely recover tens of thousands of smaller-sized phage genomes in a single study ^4,62^, huge phages remain underrepresented. We expect HPGC to become a robust resource for identifying huge phages in metagenomes, especially when combined with manual genome curation and/or single-molecule sequencing ^63^, which could partly resolve the fragmentation issues that emerge during short-read-based assembly and hamper the identification and annotation of huge phage genomes.

With the fast growth of sequencing data in public repositories, tools that can deal with the large datasets accurately and efficiently are of paramount importance. Pebblescout ^38^, Logan ^41^, Serratus ^64^, and BIGSI ^65^ are among such tools that substantially facilitate access, search, and scalable mining of the massive and largely untapped metagenome sequencing data. In this study, we showed how ecological analyses of a viral clade could be extended by utilizing some of these tools, leading to the discovery of abundant but overlooked viral clades (Figures 1d and 4).

In the case of Jug phages, we found that some essential viral proteins, such as TerL and MCP, are highly similar among group members (Figure 2), which enables the reliable identification of Jug phages in metagenomic assemblies. The expanded genomic diversity via Logan-assembly revealed the wide distribution of Jug phages and likely frequent cross-animal-host transmission (Figure 4). The transmission of the Jug phages between infant and mother, and among healthy individuals and patients via FMT (Figure 5a) suggests that Jug phages can readily spread in the human population, explaining their wide distribution (Figure 4). In addition, we identified relatives of Jug phages in the gut of ruminants (Figure 2a). Notably, in each of the analyzed gut metagenomes, with a single exception, we detected only one Jug phage genome type. It seems likely that Jug phages possess superinfection exclusion mechanisms that prevent infection of the host bacteria by other genome types. However, the possibility cannot be ruled out that we failed to retrieve Jug phage genomes from samples containing multiple genome types because the assemblies could be highly fragmented due to the high nucleotide identity between genomes from the same group.

The pronounced discrepancy between the high overall genome similarity and the low TerL identity among the Alpha, Gamma, and Delta groups suggested that Jug phages have undergone targeted replacement of the *terL* gene via HGT (Figure 2a, Supplementary Figure 11). To our knowledge, this represents the first case of *terL* replacement in huge phages, although similar observations have been reported for crAss-like phages ^13^ and other groups of phages from the human gut ^17^. This finding challenges the common practice of using TerL as a universal phylogenetic marker for Caudovirales ^66^. The *terL* replacement might reflect adaptive fine-tuning of the DNA packaging mechanisms in response to ecological pressures or host-specific constraints, or could be a route of escape from host defenses.

In contrast to crAss-like phages ^13^, Lak phages ^35^, and other gut phages ^67^, Jug phages show no evidence of using alternative genetic codes. However, similar to crAss-like phages ^13^, gene fragmentation through insertion of group I self-splicing introns and inteins is a common feature of Jug phage genomes (Figures 2a and d). Given that many viral genes remain uncharacterized, fragmentation obscures gene prediction and functional inference. Integrating metatranscriptomic sequencing and reads mapping with structure-based protein annotation could help resolve gene boundaries and assess whether fragmented ORFs are spliced or co-translated (Figure 2d).

Jug phages were predicted to infect *Bacteroides* and/or *Phocaeicola*, which are dominant in human gut microbiotas and play essential roles in degrading dietary and host-derived glycans, thereby contributing to short-chain fatty acid production and gut health ^68,69^. The increased occurrence of Jug phages in individuals on high-fiber diets supports this predicted host–phage association (Supplementary Figure 24).

Finally, we report here the identification of calcium-translocating P-type ATPases in Jug phages and other huge phages, as well as a few phages with smaller genomes infecting Bacteroidaceae and Bacillota (Supplementary Figure 29), and show that this gene is expressed at a high level in at least some Jug phages (Figures 6e and f). These membrane proteins likely manipulate host intracellular Ca^2+^ levels, influencing host metabolism, stress response, and short-chain fatty acid production. We hypothesize that this phage-encoded calcium export system confers a competitive advantage by remodeling the host cytoplasmic environment to optimize phage replication, and/or indirectly reshape gut microbial ecology and host physiology by disrupting calcium-regulated commensal bacterial functions. Such mechanisms reflect a broader strategy in which phages deploy auxiliary metabolic functions to manipulate host cell physiology for their benefit.

### Concluding remarks

Given the broad distribution, high transcriptional activity, and diverse auxiliary functions of Jug phages, we propose that they play an active role in modulating gut microbiome composition and host physiology. Their ecological influence could stem from lysis of bacterial hosts that are major microbiome components, and/or through metabolic modulation via the activity of phage genes such as the calcium-translocating P-type ATPases. Further research is warranted to isolate Jug phages in culture, characterize their infection cycles, and experimentally validate their impact on host cells and microbial communities, as suggested by the in-depth bioinformatic analyses presented in this study.

## Methods

### Viral genome collection

Published or public potential huge phage genomes were collected from NCBI Genbank and publications via the downloading links or accession numbers if provided, as of September 30, 2024.

The NCBI phage genomes with a minimum length of 200 kbp were collected with the keyword “phage”, and the “Sequence length” was set from “200000” to “1000000”, which retrieved a total of 954 genomes (“NCBI_phage_200kbp”). To avoid missing any genomes of huge phages, which are somehow named “virus” in NCBI, we also searched “virus” with the same length range, and retrieved a total of 943 genomes (“NCBI_virus_200kbp”). Local genome comparison indicated that all the previously reported isolated huge phages ^27^ were included in the collected NCBI sequences.

The viral datasets published before August 31, 2024, were downloaded according to the descriptions in the corresponding publications (if available). A detailed description of the publications is provided in Supplementary Table 1. The viral genomes were filtered to retain those with a minimum length of 200 kbp. We also retrieved some metagenomic datasets sequenced with third-generation sequencing technology, such as Nanopore. For example, the Nanopore sequences (passed default Guppy quality control) reported recently were provided by the authors ^70^; those with a minimum length of 200 kbp were screened. The IMG/VR v4 database was filtered to retain the viral genomes (1) with a minimum length of 200 kbp, (2) not assigned as “Megaviricetes” (via the corresponding IMG/VR mapping file), and (3) with the corresponding sample is not under the JGI Data Utilization Status of “Restricted”, which obtained a total of 5,181 sequences.

### Huge phage genome identification

The collected viral sequences were first evaluated by geNomad (parameter: end-to-end) ^60^. The geNomad results were parsed using the “Conservative standard”, i.e., virus_score ≥ 0.80, n_hallmarks ≥ 1, mark_enrichment ≥ 1.5, and fdr ≤ 0.05. The geNomad hits were then evaluated by VirSorter2 ^71^ with the following parameters: --keep-original-seq, --include-groups “dsDNAphage,NCLDV,RNA,ssDNA,lavidaviridae”. And only the sequences with a VirSorter2 score of ≥ 0.5 were retained for further analyses; otherwise, they would be excluded.

Then, CheckV ^59^ (parameter: end_to_end) was used to analyze the virus-containing sequences to remove the host regions. Subsequently, the output files of “proviruses.fna” and “viruses.fna” from CheckV were combined (termed as “checkv.fa”). The CheckV provirus sequences were re-run in a second CheckV analysis to obtain the relevant information on the virus fragments.

The “checkv.fa” sequences (pure viral fragments) are evaluated by another run of VirSorter2 analyses with the parameters as follows: “--seqname-suffix-off --viral-gene-enrich-off --provirus-off --prep-for-dramv --include-groups dsDNAphage,NCLDV,RNA,ssDNA,lavidaviridae”. Subsequently, the genomes were filtered based on information from the second CheckV and the second VirSorter2 analyses to retain those with (1) viral_gene >0 (i.e., Keep1), or (2) viral_gene =0 and (host_gene = 0 or score >=0.95 or hallmark >2) (i.e., Keep2), following a widely used online procedure ^72^.

The retained sequences were subjected to a BLAST search to check for replication issues that may arise in the *de novo* assembly. If a given sequence’s first half and second half shared at least 50% of their length with ≥ 99% nucleotide sequence similarity, it was flagged, and a manual check was performed to exclude any assembly issues.

In the final step, if a given sequence was assigned as “NCLDV” by geNomad while as “Caudoviricetes” by Virsorter2, or vice versa, then a manual check was performed for confirmation based on the taxonomic information (most similar UniProt BLASTp hits) of all the protein-coding genes of the sequence.

### Genome clustering

To remove redundant genome sequences that may be reported from different studies or different assemblies, all the identified huge phage genomes were clustered using the rapid genome clustering approach provided by CheckV (https://bitbucket.org/berkeleylab/checkv/src/master/) ^59^, with the parameters set as “min_ani = 100, min_tcov = 100”, to obtain the non-redundant genome set. These non-redundant genomes were further clustered into species-level clusters using the same approaches, with the parameters of “min_ani = 95, min_tcov = 85”.

### Manual curation of the Jug phage genomes

To curate the Jug phage genomes, the corresponding paired-end read files were downloaded from NCBI and assembled using metaSPAdes version 3.15.2 ^73^ with the kmer set of “21,33,55,77,99”. The assembled scaffolds were then searched against the HPGC Jug phage genomes using BLASTn for Jug phage-related scaffolds. The target scaffolds were manually curated via read mapping, scaffold extension, assembly error fixation, and reassembly, as previously described ^74^, with step-by-step procedures available at https://ggkbase-help.berkeley.edu/genome_curation/scaffold-extension-and-gap-closing/. The scaffold extension is heavily based on the unplaced paired-end reads. For example, for a paired-end read_1 and read_2, if read_1 is well-mapped to the reference while read_2 is not, then read_2 is named the unplaced paired-end reads. When performing the reads mapping using Bowtie2 or other alternative tools, we usually only output the mapped reads to the SAM/BAM files to save computational storage space. This could be conducted by adding the parameter of “--no-unal”. However, this will exclude both read_1 and read_2 in the abovementioned example. The shrinksam (available at https://github.com/bcthomas/shrinksam) was designed to output both reads, even if only one is well mapped. Alternatively, an in-house script could be developed to this end (for example, using the pysam tool that is available at https://github.com/pysam-developers/pysam). Please note that manual curation is time-consuming and should be performed only for those that are of high interest.

### Protein-coding gene prediction and annotation

The protein-coding genes from each of the 139 curated Jug phages were predicted using Prodigal version 2.6.3 (single mode) with code 11 ^75^. The functional domains of the Jug phage proteins were predicted using hmmsearch from HMMER version 3.3.2 ^76^ against the Pfam database ^77^. The results were parsed using cath-resolve-hits version 0.16.10-0-g99edb28 ^78^, then the identification of a domain in a protein was filtered by an independent e-value (“indp-evalue”) threshold of 1e-5.

For protein structure prediction, the protein sequences were first clustered using CD-HIT with the parameters of “-c = 1, -aS = 1”. For representatives of each cluster, protein structure was predicted using ColabFold (downloaded on November 22, 2024) ^79,80^, with the following parameters: “--num-recycle 3, --use-gpu-relax, --amber, --stop-at-score 70”. For structure annotation, the predicted structures with a pLDDT score of ≥70 were searched against the protein data bank (PDB) ^81^ and the predicted structures of BFVD ^82^, using FoldSeek 9.427df8a ^83^ with the “easy-search” command (−c = 0.5, --cov-mode = 0). Only those hits with a TM score ≥0.5 (“alntmscore”) were retained for further analyses. The structure prediction and annotation of specific proteins, such as large terminase, major capsid proteins, and DNA polymerase, were confirmed via online FoldSeek search ^83^ at https://search.foldseek.com/search.

### Identification of group I introns

To globally identify potential group I introns in the Jug phage genomes, we downloaded the CM format files of all the group I intron subtypes reported recently ^84^ and searched against the Jug phage genomes using cmscan with the INFERNAL version 1.1.4 ^85^ with an e-value threshold of 1e-4. If a specific region was predicted with multiple subtype hits, only the one with the lowest e-value was retained and counted.

### Protein similarity comparison of different groups of Jug phages

To identify the single-copy protein clusters shared by all four groups, the predicted protein sequences were clustered using CD-HIT version 4.8.1 ^86^ with ≥70% identity (−c 0.7 -aS 0.9 -G 0), and the single-copy protein clusters were identified accordingly. We screened all the single-copy protein clusters, allowing a given cluster to be present in at least half of the genomes in each group (Supplementary Table 7).

Because the genomes of groups Alpha, Gamma, and Delta shared high genome-wide similarity while having low TerL identity, we investigated the single-copy protein clusters shared by all three groups for other lower similarity ones. The predicted proteins of groups Alpha, Gamma, and Delta were clustered using CD-HIT version 4.8.1 ^86^ with ≥70% identity (−c 0.7 -aS 0.9 -G 0). For each of the clusters, including members from Alpha, Gamma, and Delta, the average similarity of members against the representative was calculated (Supplementary Table 5).

As the genomes of groups Alpha and Beta shared high *TerL* identity but low genome-wide similarity, we investigated the single-copy protein clusters shared by both groups for other highly similar ones. The predicted proteins of group Alpha and Beta were clustered using CD-HIT version 4.8.1 ^86^ with ≥70% identity (−c 0.7 -aS 0.9 -G 0). For each of the clusters with ≥5 members from group Alpha and ≥5 members from group Beta, if the cluster representative was from group Alpha, the average similarity of group Beta members against it was calculated; otherwise, the average similarity of group Alpha members against it was calculated (Supplementary Table 4).

### Large terminase identity within other HPGC genome clusters

The other HPGC huge phage genomes (excluding Jug phage ones) are clustered at 90% nucleotide sequence similarity across at least 50% of the genome length. A total of 212 clusters were identified with 5 or more members. The large terminase proteins were predicted using geNomad version 1.5.2 ^60^. Among the 79 clusters with at least 2 identified large terminase proteins, their within-cluster large terminase protein identity was determined using BLASTp.

### Phylogenetic analyses of Jug phages based on TerL

The TerL fragments from curated Jug phage genomes were first concatenated into one piece and aligned with the full-length TerLs of Jug phages, the TerLs of Jug phage relatives, and the TerLs from the gut of sheep and water deer. All the HPGC proteins (excluding those from Jug phages) were searched against the Jug phage TerL sequences, and those hits with an e-value of ≤1e-5, ≥30% identity, and a minimum length of 400 aa were determined as Jug phage relatives. For references, we also included NCBI Genbank TerLs with a minimum identity of 20% to those of Jug phages. The alignment of TerL sequences was performed using MUSCLE v3.8.31 ^87^ and filtered using trimAl v1.4.rev22 ^88^ to remove those columns comprising ≥90% gaps. The phylogenetic trees were constructed using IQ-TREE ^89^ with the parameter “-bb 1000, -m LG+G4”.

### Bacterial host prediction

We utilized both iPhoP version 1.3.3 ^90^ and local CRISPR-Cas spacer targeting to predict the host of Jug phages. The iPhoP analysis was performed using default parameters. For local CRISPR-Cas spacer targeting analyses, we firstly predicted the repeat regions from all the metagenomic assembled scaffolds (from the samples we used for manual curation of Jug phage genomes) using PILER-CR version 1.06 ^91^ with default parameters. The spacers were extracted from the predicted repeat regions with Cas genes presented within 1000 bp. The curated Jug phage genomes were searched against the spacers with an e-value threshold of 1e-4, and only those spacers matched with ≤1 mismatch across the whole spacer length were counted as targeting spacers. The Cas genes were predicted by an HMM search of the predicted protein-coding genes of repeat-containing scaffolds against an in-house Cas reference database that was primarily from the TIGRFAM dataset ^92^. The taxonomic assignment of the scaffolds with targeting spacers was conducted by comparing their protein-coding genes against the UniprotKB database ^93^ using MMseqs2 version 15.6f452 ^94^.

### Comparison against the crAss-like phage and Lak phage genomes

To compare the Jug phage genomes against crAssphage (145-192 kbp) and Lak phages (408-660 kbp), their genomes were downloaded from the corresponding publications ^13,35,36^ and searched using BLASTn with an e-value threshold of 1e-5 and a minimum alignment length of 1000 bp. The protein-coding genes of crAss-like phages and Lak phages were downloaded from the corresponding publications and compared against the large terminase and major capsid protein sequences of Jug phages using BLASTp with an e-value threshold of 1e-5.

### The co-occurrence analyses of Jug phages and predicted bacterial hosts

To evaluate the distribution of Jug phages, crAssphage, Lak phages, and pBI143, the quality paired-end reads from 125 samples were mapped to the genomes using Bowtie2 version 2.4.4 ^95^ with default parameters. The Shriksam tool (https://github.com/bcthomas/shrinksam) was used to exclude unmapped pairs from the SAM file, which was subsequently sorted and converted to the BAM format using Samtools ^96^. The number of mapped reads to each genome across samples was generated by CoverM version 0.7.0 (“contig” mode) (https://github.com/wwood/CoverM) using the “count” method with the parameters set as “--min-read-aligned-percent 95 --min-read-percent-identity 90 --contig-end-exclusion 75 -m count”.

The relative abundance of the predicted bacterial host across the 125 metagenomic samples was determined using ribosomal protein S3 (rpS3). First, the protein-coding genes from all the assembled scaffolds were predicted using Prodigal version 2.6.3 (meta mode) with code 11 ^75^. Then, the rpS3-coding genes were predicted using hmmsearch from HMMER version 3.3.2 ^76^ against the kofam ^97^ by identifying those matching K02982 (score threshold = 108.13) or K02984 (score threshold = 67.70). The taxonomic assignment of the identified rpS3 was conducted by searching against the rpS3 database retrieved from GTDB (gtdb_r226) ^98^ using BLASTp. For each sample, to calculate the relative abundance of the predicted bacterial hosts, the sequencing coverage of the corresponding rpS3-encoding scaffold was divided by the total sequencing coverage of all rpS3-encoding scaffolds in the sample.

### Pebblescout and Logan search

The curated Jug phage genomes were individually used for an online Pebblescout search at https://pebblescout.ncbi.nlm.nih.gov/ with the target database as of “Metagenomic” ^38^. The information on all the targeted metagenomic samples was downloaded for further analysis. The metagenomic samples with a Pebblescout “%coverage” of ≥70 were selected for *de novo* assembly and manual genome curation for Jug phage genomes.

The first 1000 nt (from 5’ end) of the nucleotide sequences of the Jug phage *TerL* genes were used for online Logan search at https://logan-search.org/dashboard with a threshold of ≥0.7 and with the “Group” parameter as “All” ^41^. The contigs of the targeted metagenomic samples were downloaded via “*aws s3 cp s3://logan-pub/c/*{*sample*}*/*{*sample*}.*contigs*.*fa*.*zst*. *--no-sign-request*”, in which “{sample}” represents the NCBI SRA ID of the corresponding sample, which was obtained from the Logan search. The downloaded contigs with a minimum length of 1000 bp were searched against the curated Jug phage genomes using BLASTp, and those with ≥90% similarity across ≥90% of their length were considered as reliable hits. The protein-coding genes were predicted from all the retrieved contigs ≥1000 bp using Prodigal version 2.6.3 ^75^ using the parameters of “-m -p meta”, and searched against the *TerLs, rpoC*, and the 38 single-copy protein clusters from the 139 curated Jug phage genomes using BLASTp.

### Protein clustering analyses of curated and Logan-retrieved Jug phage genomes

The protein-coding genes were predicted from all the manually curated genomes and the Logan-retrieved Jug phage genomes using Prodigal version 2.6.3 ^75^ using the parameters of “-m -p meta”. The protein sequences were clustered using CD-HIT version 4.8.1 ^86^ using the parameters of “-c 0.7 -aS 0.9 -G 0”, which meant ≥70% identity across ≥90% length of both the query and target sequences. The distribution and sharing of each protein cluster among different groups, or different animal hosts of each group, were evaluated. If at least one genome of a given group or host animal had a protein in a given protein cluster, then the corresponding protein cluster was considered to be present in the group or host animal. The annotation of the proteins was performed as described above.

### Analysis of Jug phages during fecal microbiota transplantation

The Logan-retrieved genomes included samples from fecal microbiota transplantation (FMT), which were used to investigate the transmission of Jug phages among humans. In the study by Smith et al ^45^, the authors treated 22 patients with mild to moderate ulcerative colitis by FMT from two healthy donors. The FMT was conducted by capsules (CAPS) or enema (ENMA) for maintenance dosing. For comparison, 11 of the patients received antibiotic pretreatment before the FMT. The FMT was performed 6 times, followed by three follow-up evaluations conducted in 2 weeks, 6 weeks, and 14 weeks (unless otherwise stated) after the last maintenance dose. The fecal samples were collected for metagenomic analysis at the beginning, before antibiotic pretreatment, before each maintenance dosing, and before each follow-up evaluation. Thus, 10 (without antibiotic pretreatment) or 11 (with antibiotic pretreatment) metagenomic samples were generally available for each recipient, unless some samples were not sequences for various reasons.

The raw paired-end metagenomic reads from all donors and recipients of all time points and filtered using fastp version 0.22.0 ^99^ with default parameters. The quality reads were assembled using metaSPAdes version 3.15.2 ^73^ with the kmer set of “21,33,55,77,99”. The assembled scaffolds were searched against the curated Jug phage genomes using BLASTn, and the confirmed Jug phage-related scaffolds were manually curated. The quality reads of each recipient at each time point were mapped to the curated Jug phage genome of the donor with Bowtie2 version 2.4.4 ^95^ with default parameters. The sequencing coverage of the Jug phage at each time point was calculated by CoverM version 0.7.0 (“contig” mode) (https://github.com/wwood/CoverM) using the “trimmed_mean” method with the parameters set as “--min-read-aligned-percent 99 --min-read-percent-identity 100 --min-covered-fraction 75 --contig-end-exclusion 75 -m trimmed_mean”.

### Analysis of Jug phages in the guts of humans with dietary intervention

In the study reported by Delannoy-Bruno et al, 12 and 14 overweight or obese men or women completed dietary intervention studies 1 and 2, respectively ^46^. In study 1, the gut microbiome of the participants contained ≥0.1% of *Phocaeicola vulgatus* and/or ≥0.1% of *Bacteroides thetaiotaomicron, B. cellulosilyticus, B. uniformis*, or *B. ovatus*. Each of the participants was asked to maintain their usual diet for 4 days, on days 5 to 14, they were asked to consume only HiSF-LoFV meals. Starting from day 15, the participants were supplemented with pea fibre-containing snacks while continuing the HiSF-LoFV meals. The snack was taken once per day (i.e., one bar per day) on days 15 and 16, twice per day on days 17 and 18, then three times per day from day 19 through day 35. Then the participants returned to HiSF-LoFV meals without a snack bar for another 14 days. In study 2, the participants were not prescreened for the presence of *Bacteroides* species. The participants were asked to consume HiSF-LoFV meals from days 2-11, then with a 2-fibre snack prototype once on day 13, twice on day 14, and three times on days 14-25. On days 26-35, the participants consumed only HiSF-LoFV meals without fibre snacks. Afterward, they were supplemented with a 4-fibre snack prototype once on day 36, twice on day 37, and three times per day on days 38-49.

We identified Jug phages using the Logan assembly in three of the participants (i.e., S11, S13, and S15), with Jug phages detected in S13 samples of both studies, in S11 samples of study 1(completed both studies), and in S15 samples of study 1 (completed study 1 only). The quality control of raw metagenomic reads, assembly, and manual genome curation was performed as for the FMT samples. The identification of *Bacteroides* and *Phocaeicola* species in the samples was based on rpS3 as described above. For each participant, the unique rpS3-encoding scaffolds were used as read-mapping reference along with the corresponding curated Jug phage genome, then the quality paired-end reads of each time point were mapped using Bowtie2 version 2.4.4 ^95^ with default parameters. The sequencing coverage of the Jug phage at each time point was calculated by CoverM version 0.7.0 (“contig” mode) (https://github.com/wwood/CoverM) using the “trimmed_mean” method with the parameters set as “--min-read-aligned-percent 99 --min-read-percent-identity 100 --min-covered-fraction 75 --contig-end-exclusion 75 -m trimmed_mean”. The taxonomic assignment of the identified rpS3 was conducted by searching against the rpS3 database retrieved from GTDB (gtdb_r226) ^98^ using BLASTp. For each sample of each participant, the relative abundance of the *Bacteroides* and *Phocaeicola* species was calculated as the sequencing coverage of the corresponding rpS3-encoding scaffold divided by the total sequencing coverage of all rpS3-encoding scaffolds in the sample. To make it comparable, the presence ratio of each Jug phage in each sample was calculated as the sequencing coverage of the Jug phage genome divided by the total sequencing coverage of all rpS3-encoding scaffolds in the corresponding sample.

### Analysis of Jug phages for transcriptional activity

Pebblescout search ^38^ was used to find available metatranscriptional datasets with Jug phages. The Jug phage genomes from three adult men, each with four DNA and four RNA samples ^40^, were reconstructed via *de novo* assembly and manual curation as described above. To evaluate *in situ* transcriptional activities in the human gut, the RNA reads were mapped to the corresponding Jug phage genomes using Bowtie2 version 2.4.4 ^95^, the alignment was filtered using CoverM version 0.7.0 (“contig” mode) using the “count” method with the parameters set as “--min-read-aligned-percent 99 --min-read-percent-identity 100 --contig-end-exclusion 75 -m count”. The protein-coding genes were predicted using Prodigal version 2.6.3 (single mode) ^75^, and RPKM (reads per kilobase per million mapped reads) of each protein-coding gene was calculated. The annotation of the proteins was performed as described above.

### Analyses of TerD domain-containing proteins

To reveal the distribution of *TerD*-domain containing proteins in the Unified Human Gastrointestinal Genome (UHGG) database, the unique protein sequences (“uhgp-100.tar.gz”) were downloaded from http://ftp.ebi.ac.uk/pub/databases/metagenomics/mgnify_genomes/human-gut/v2.0/protein_catalogue. The proteins were first searched against the Pfam *TerD* domain HMM database (i.e., PF02342_23) using hmmsearch from HMMER version 3.3.2 ^76^, and parsed with cath-resolve-hits version 0.16.10-0-g99edb28 ^78^. The proteins with an indp-evalue ≤1e-5 were searched against the whole Pfam HMM database, and parsed using the cath-resolve-hits version. For a given protein, if the indp-evalue targeting another Pfam domain was lower than that of the *TerD* domain, then the protein was believed to have no *TerD* domain. The identified *TerD*-domain containing proteins were assigned to the genome with taxonomy based on the associated UHGG metadata (“genomes-all_metadata.tsv”). Notably, we found that the UHGG protein database contained many Jug phage proteins via BLASTp search (with >99% identity). For example, 236 out of the 427 proteins from MGYG000275746 (UHGG assigned taxonomy: d_ Bacteria;p_Firmicutes_A;c_Clostridia;o_TANB77;f_CAG-508;g _CAG-245;s_ CAG-245 sp000435175) were actually Jug phage proteins. The *TerD*-domain containing proteins matching those from Jug phages were excluded when performing the genome and taxonomy assignment.

### The analyses of calcium-translocating P-type ATPase

The calcium-translocating P-type ATPase genes (ATPase hereafter for short) identified in Jug phages according to the Pfam domain search usually contained four conserved Pfam domains: “Cation_ATPase_N”, “E1-E2_ATPase”, “Cation_ATPase”, and “Cation_ATPase_C”. The annotations were confirmed by protein structure prediction using ColabFold (downloaded on November 22, 2024) ^79,80^ and compared using FoldSeek 9.427df8a ^83^ with the “easy-search” command (−c = 0.5, --cov-mode = 0).

The scaffolds assembled from the 7 adult metagenomics with transcriptionally active Jug phages were conducted for protein-coding gene prediction using Prodigal version 2.6.3 (−m, -p = meta) ^75^, followed by a BLASTp search against the Jug phage ATPases with an e-value threshold of 1e-5. The proteins with a minimum length of 500 aa and ≥400 aa aligned length were considered as hits, which were predicted for domains using hmmsearch from HMMER version 3.3.2 ^76^ against the Pfam version 37 database ^77^. Those proteins with two of the four domains were considered ATPases. All those predicted ATPases from the 7 metagenomic samples and those from curated Jug phages were clustered at ≥99% identity (−c 0.99 -aS 0.9 -G 0) using CD-HIT version 4.8.1 ^86^, and the representatives were used for phylogenetic tree analyses. The taxonomic assignment of the ATPases was determined based on the proteins encoded on the corresponding scaffolds, and confirmed according to the phylogeny given the truth that those from phylogenetic relatives were usually clustered together on the tree. The identification of the calcium-translocating P-type ATPases from NCBI Genbank and other HPGC phage genomes was conducted using the same pipeline.

Structural annotation of calcium-transporting P-type ATPases was performed using a combination of InterProScan and NCBI CDD analyses. Functional domains, namely the actuator (A), phosphorylation (P), and nucleotide-binding (N) domains, were defined based on conserved features, although the exact residue ranges vary across different proteins. The transmembrane helices were predicted using the DeepTMHMM web server (https://dtu.biolib.com/DeepTMHMM) and grouped into three representative membrane-spanning regions (M1–2, M3–4, M6–10) for visualization and comparison. All domains were visualized in PyMOL version 3.0.3 ^100^ using cartoon representation and domain-specific colors.

The ATPase structures from human (PDB: 7e7s) and *Listeria monocytogenes* (LMCA1) were used as references to compare the spatial positions of the conserved “DKTGT” motif in phage-derived homologs. The Cα–Cα distance between corresponding “D” residues in reference and phage proteins was computed using PyMOL to assess spatial shifts in the phosphorylation loop. The “DKTGT” motifs were rendered as sticks for clarity. Calcium ions in reference structures were automatically identified by selecting atoms with the element CA and residue name CA. Known calcium-coordinating residues in the reference proteins were specified manually, and their distances to calcium ions were calculated. To identify corresponding residues in target proteins, all Cα atoms in the target were searched for spatial proximity to each reference residue. The nearest residues were recorded as potential calcium-coordinating candidates. Distances between these target residues and the calcium ions in the reference structure were then measured. This mapping procedure was conducted separately for both human and Listeria monocytogenes reference structures, enabling comparative evaluation across target proteins, including those derived from phages. Coordinating residues were shown as sticks, colored dirty violet in the reference and hot pink in the target proteins.

## Data availability

The genome sequences of the Huge Phage Genome Collection (HPGC), the CRISPR-Cas spacer sequences used for bacterial host prediction of Jug phages, and the curated genome sequences of Jug phages are available at https://figshare.com/projects/Mac_phages_in_the_gut_of_animals/254252.

## Author contributions

L.X.C. conceived this study, reconstructed the HPGC, performed all the analyses otherwise stated below, and drafted the manuscript. A.P.C. introduced the Logan-based analyses and helped with bioinformatic analyses. Y.Q. and L.X.C. performed the protein structure analysis of the Ca^2+^ ATPases. E.V.K., H.W. and Y.Z. contributed to the bioinformatics analyses. Y.D. provided suggestions on data analysis. H.L. helped with the gene transcriptional analysis. All authors revised and finalized the manuscript.

## Acknowledgements

We thank Dr. Jillian F. Banfield for commenting on the manuscript, Dr. Zhengshuang Hua for providing computational resources, and Dr. Xingxing Shen and Dr. Honglue Shi for helpful discussion. We thank Dr. Artem Babaian and the Logan team for guiding the use of the Logan search. We thank Lauren Lui for providing the quality-controlled Nanopore sequences. We thank the SuperComputing Center at the University of Science and Technology of China for its support of ColabFold analyses. L.X.C. is supported by the Research Program of the University of Science and Technology of China (KY2400000036). The work conducted by the US Department of Energy Joint Genome Institute (https://ror.org/04xm1d337) is supported by the US Department of Energy Office of Science user facilities, operated under contract no. DE-AC02-05CH11231. E.V.K. is supported by the Intramural Research Program of the National Institutes of Health of the USA.

