## Supplementary Figure for "A prevalent huge phage clade in human and animal gut microbiomes"

for

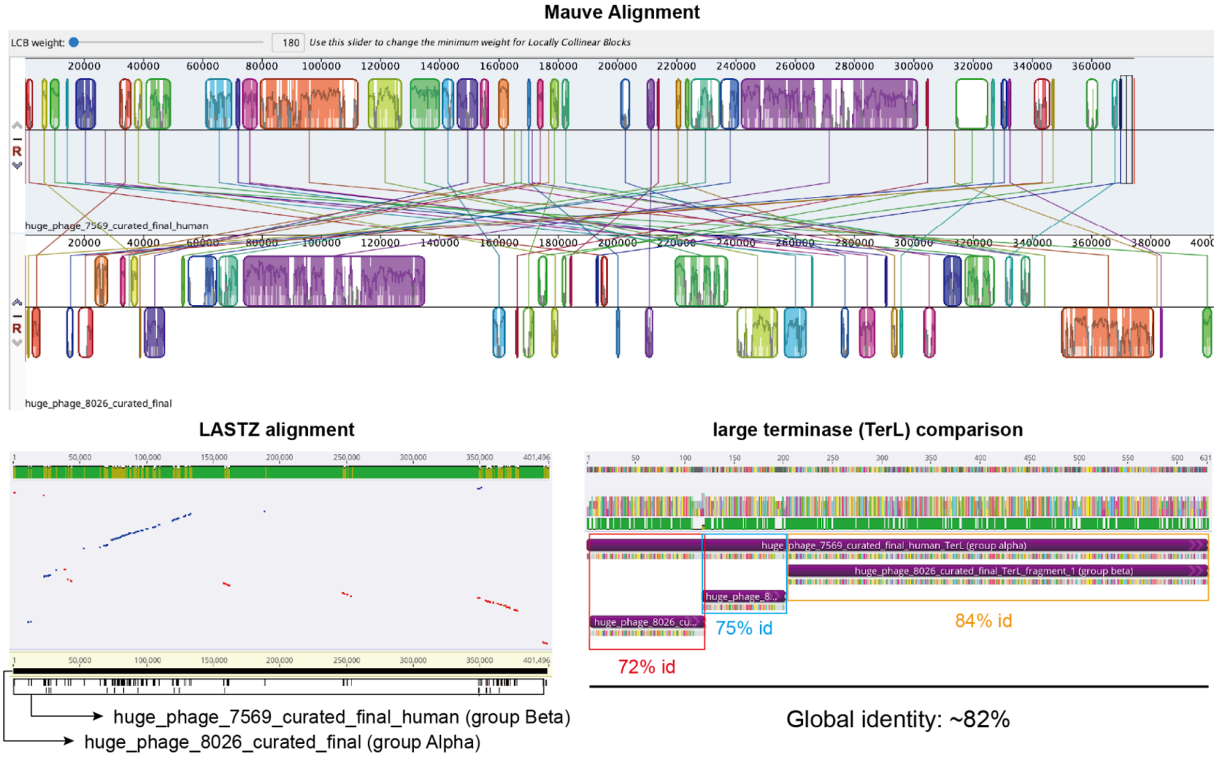


**Supplementary Figure 1 | The comparison of groups Alpha and Beta Jug phage genomes.** Jug phage genomes of huge_phage_7569 (group Beta) and huge_phage_8026 (group Alpha) were compared. The genome alignment was performed using Mauve (Darling *et al.*, 2004) and LASTZ version 1.04.15 (Harris, 2007) within Geneious Prime Build 2025-03-24 (Kearse *et al.*, 2012). The TerL alignment was conducted using the MUSCLE version 5.1 (Edgar 2004) within Geneious Prime Build, with the protein identity shown.

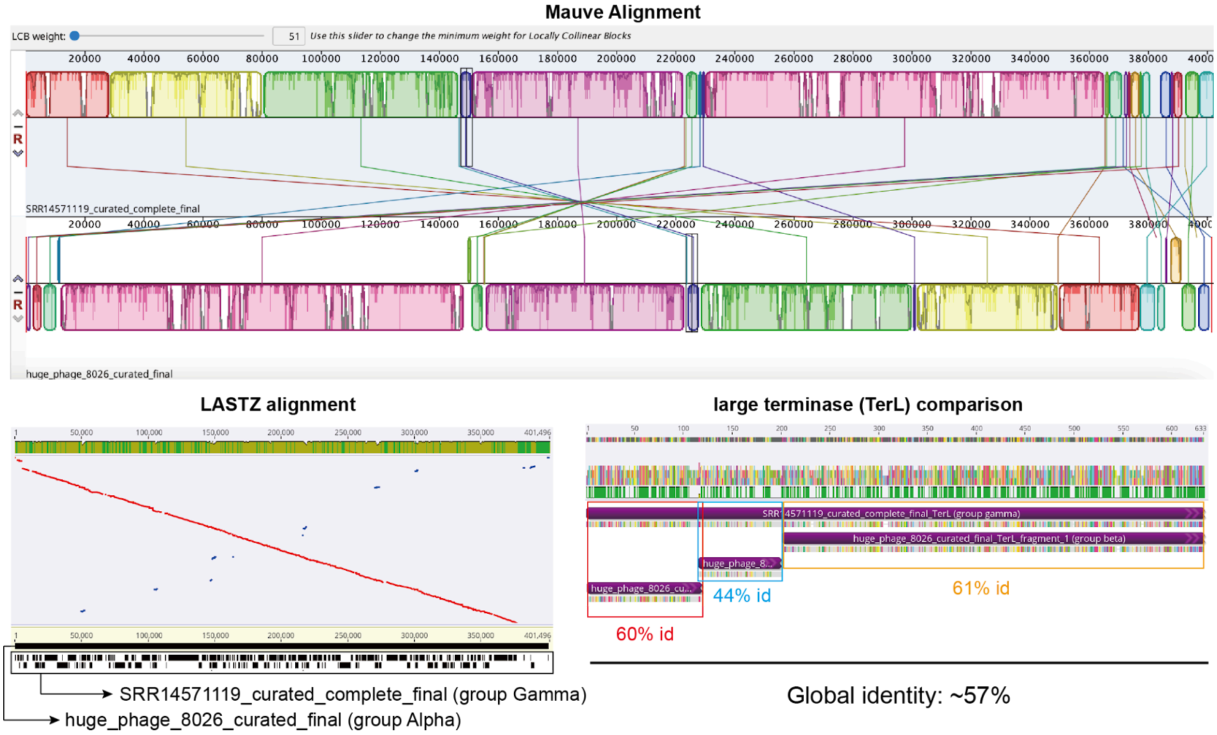


**Supplementary Figure 2 | The comparison of groups Alpha and Gamma Jug phage genomes.** Jug phages SRR14571119_curated_complete_final (group Gamma) and huge_phage_8026 (group Alpha) were compared. The genome alignment was performed using Mauve (Darling *et al.*, 2004) and LASTZ version 1.04.15 (Harris, 2007) within Geneious Prime Build 2025-03-24 (Kearse *et al.*, 2012). The TerL alignment was conducted using the MUSCLE version 5.1 (Edgar 2004) within Geneious Prime Build, with the protein identity shown.

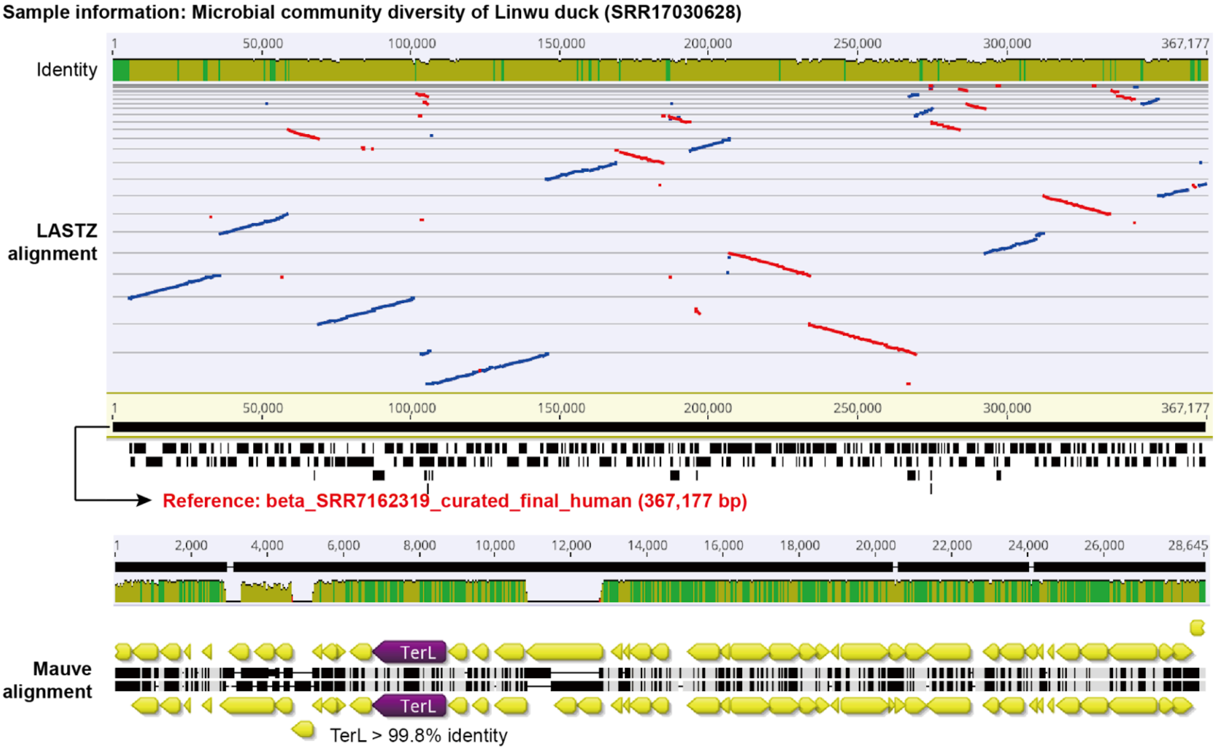


**Supplementary Figure 3 | The detection of Jug phages in the gut of ducks.** An example of group Beta Jug phage in the gut of a duck. The analysis was performed by aligning the Logan contigs against a curated Jug phage genome using LASTZ version 1.04.15 (Harris, 2007) within Geneious Prime Build 2025-03-24 (Kearse *et al.*, 2012). The Mauve alignments show the zoom-in of the large terminase (TerL) regions, with the TerL identity shown.

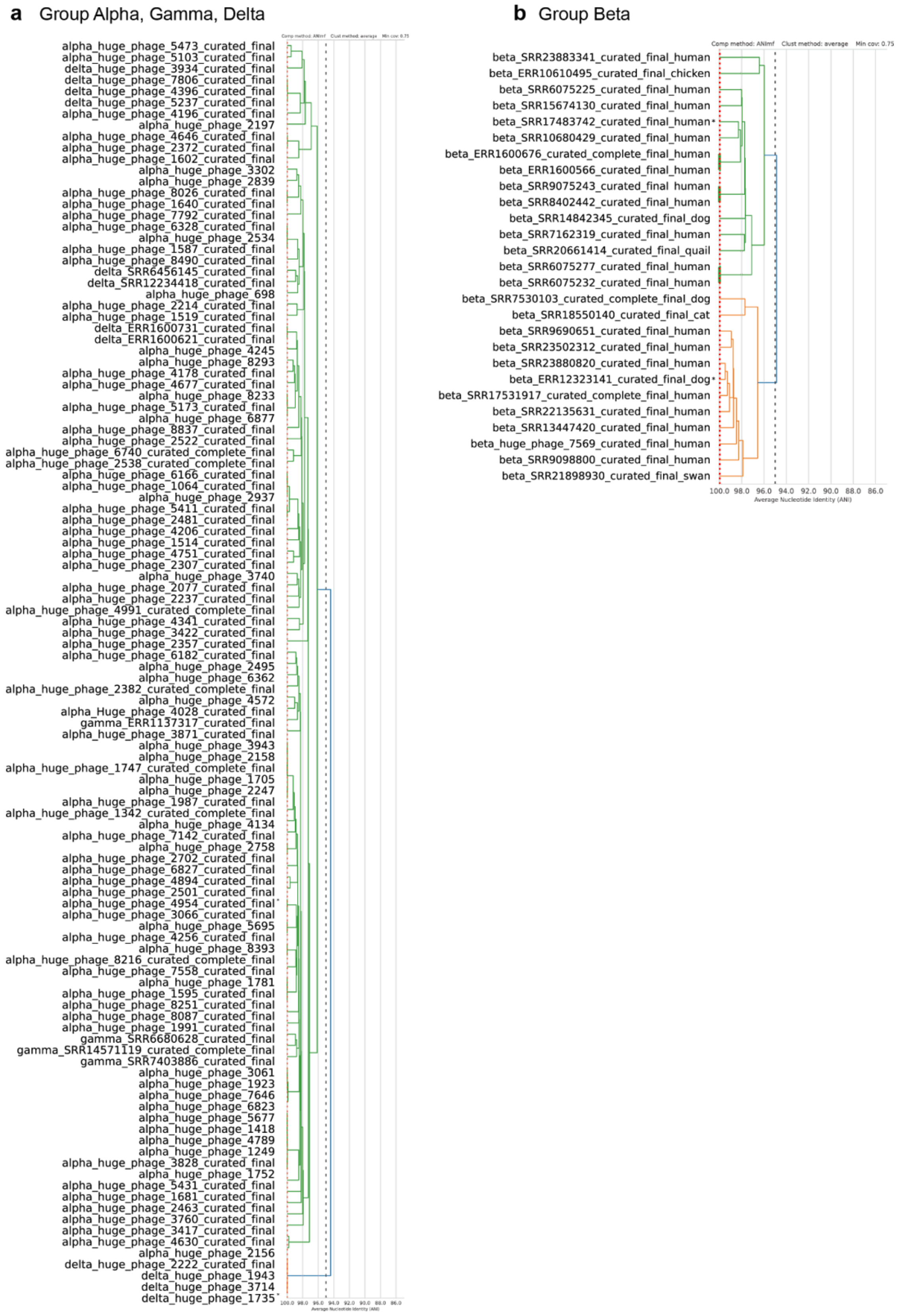


**Supplementary Figure 4 | The genome-wide clustering of the 139 curated Jug phage genomes.** (a) The genomes of group Alpha, Gamma, and Delta were clustered together, and (b) those of group Beta were clustered together. The clustering was performed using dRep version 3.2.2 (Olm *et al.*, 2017), with the parameters of "-sa 0.95 -nc 0.75".

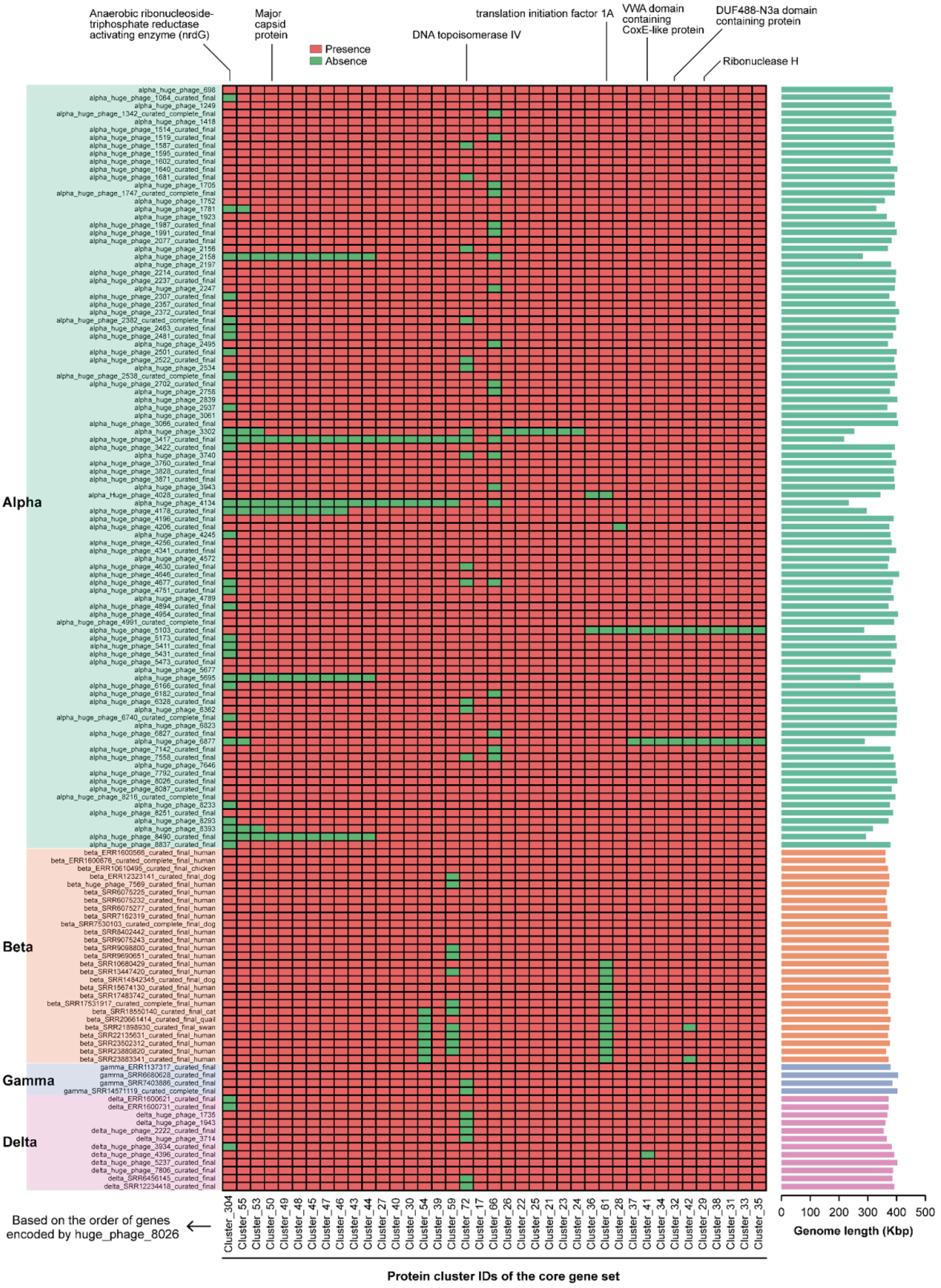


**Supplementary Figure 5 | The distribution of the 39 single-copy core gene set across the 139 curated Jug phage genomes.** The annotation of each gene is shown on the top if available; all those not shown are hypothetical proteins. The length of the genomes is shown to the right. Some of the genomes in the Alpha group lack many of the core gene set, which is likely due to their lower genome completeness as reflected by their genome length.

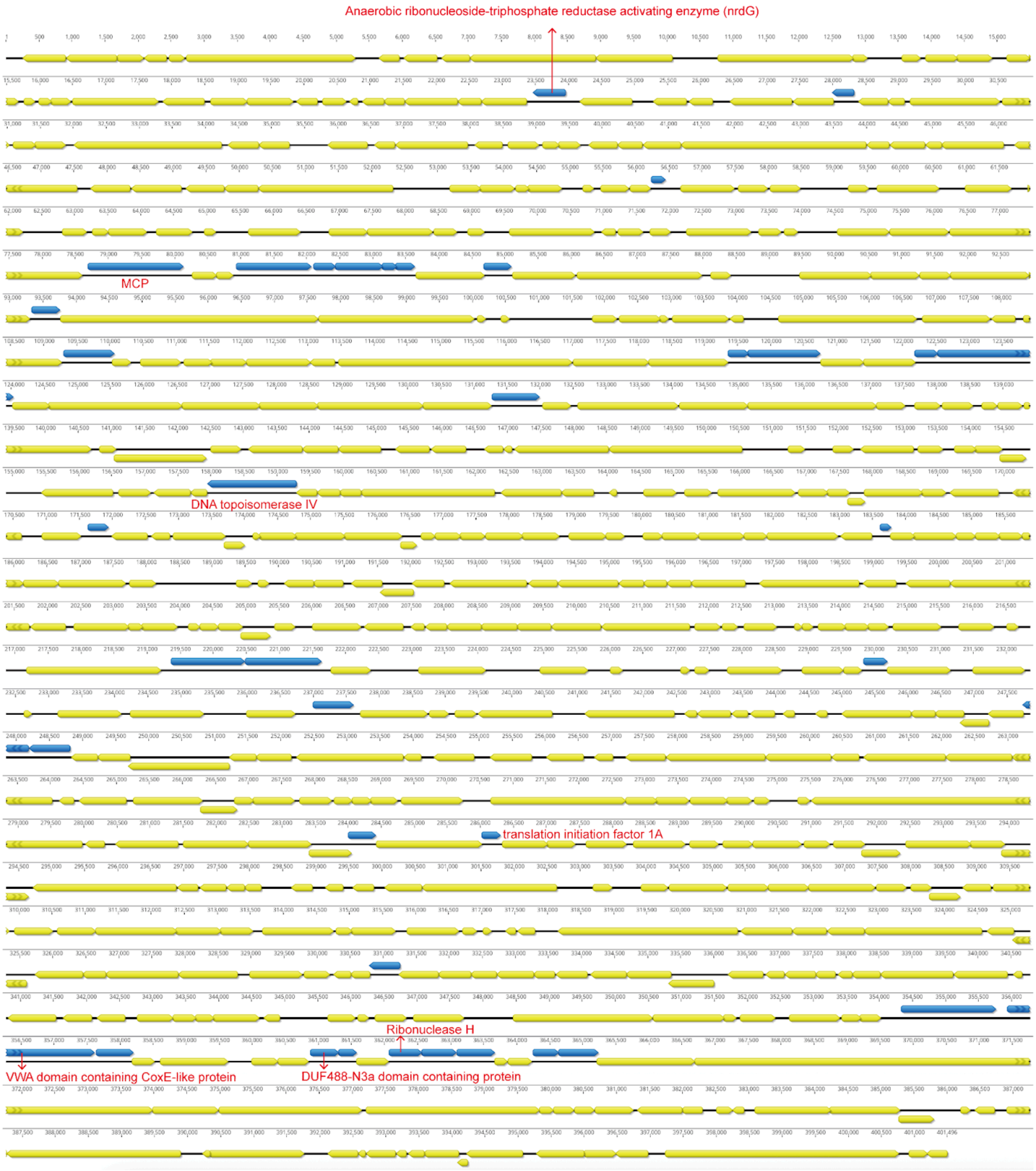


**Supplementary Figure 6 | The distribution of the 39 single-copy genes shared by all four Jug phage groups.** The protein-coding genes from all 139 curated Jug phage genomes were clustered using CD-HIT version 4.8.1 (Li and Godzik, 2006) with the parameters of "-c 0.7 -aS 0.9 -G 0". Those clusters with only one member for each genome, and including at least half of the genomes of each group, were defined as single-copy gene clusters. See Supplementary Table 7 for details of the genes. The genome of Huge_phage_8026 is used as an example to show. The genes with annotations are highlighted in red. The single-copy genes are shown in blue, and all other genes are shown in yellow. The figure was generated using Geneious Prime Build 2025-03-24 (Kearse *et al.*, 2012).

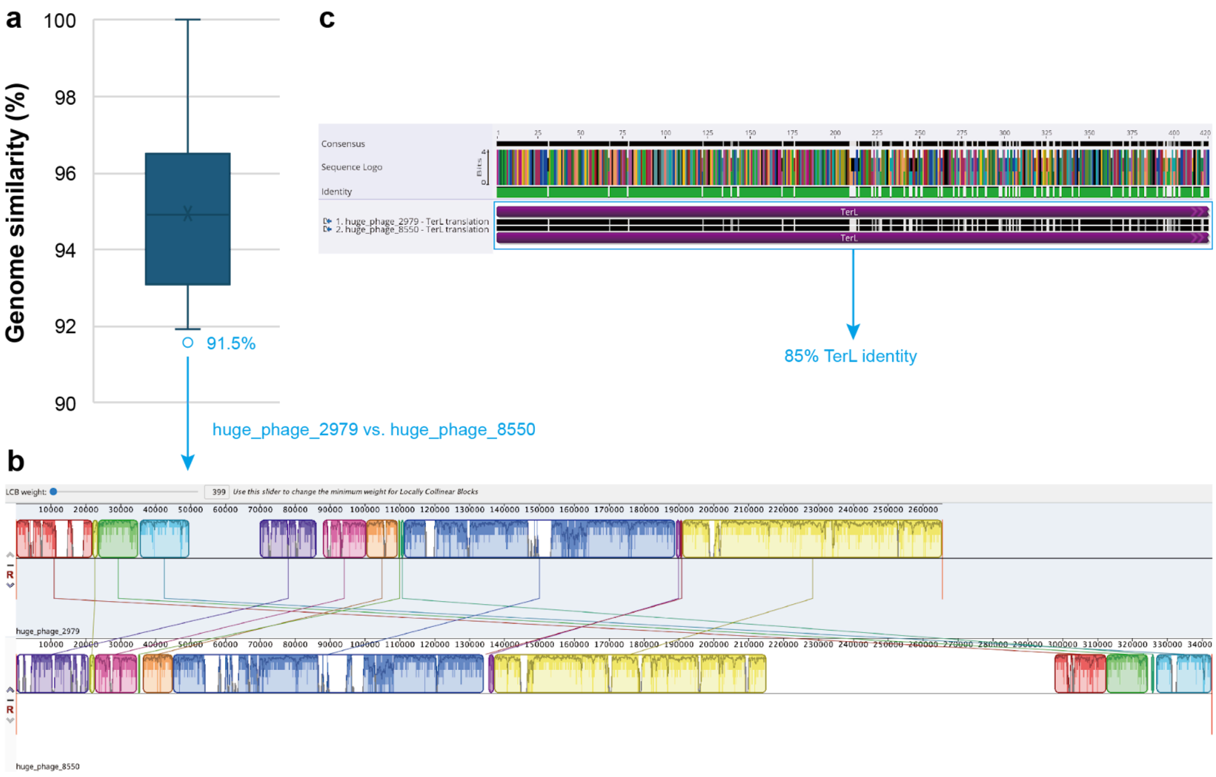


**Supplementary Figure 7 | The only other HPGC cluster had a paired TerL protein identity <90%, except for the Jug phage cluster.** The clusters were defined at ⩾90% genome-wide similarity. (a) The distribution of the paired genome-wide similarity within the cluster. (b) The whole-genome alignment of the two genomes (huge_phage_2979 and huge_phage_8550) in the cluster shared only 91.5% genome-wide similarity. The alignment was performed using Mauve (Darling *et al.*, 2004) within Geneious Prime Build 2025-03-24 (Kearse *et al.*, 2012). (c) The alignment and identity of the TerL proteins of the two genomes. The alignment was conducted using MUSCLE version 5.1 (Edgar 2004) within Geneious Prime Build 2025-03-24.

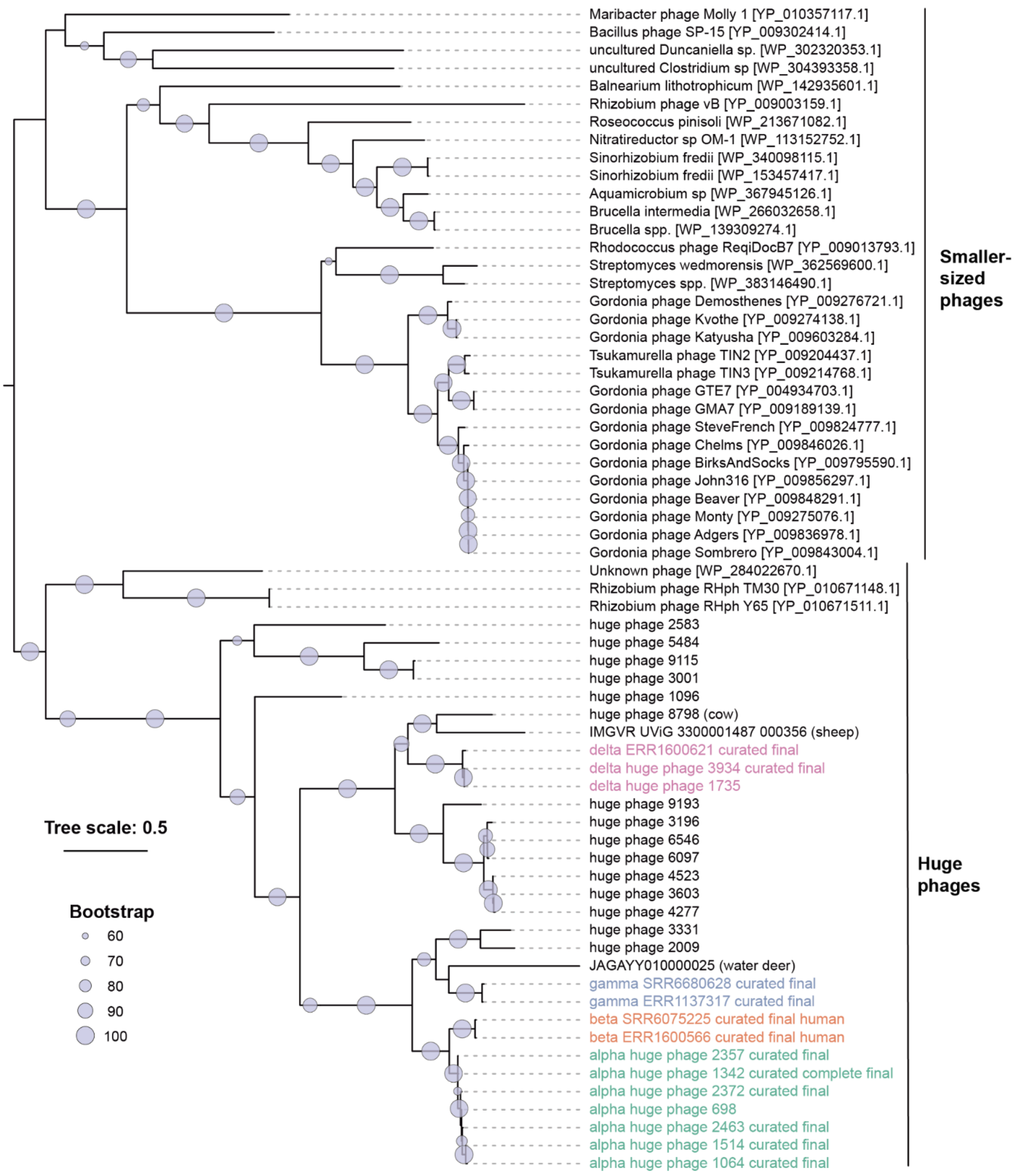


**Supplementary Figure 8 | The full phylogeny of Jug phages and related viruses based on the large terminase (TerL).** The related viruses were from NCBI by searching the TerLs of Jug phages against NCBI RefSeq and IMG/VR v4 using BLASTp. Only those hit TerLs with a minimum identity of 20% to Jug phage TerL were included (all of the identities are 20-30%). The TerL protein sequences were aligned using MUSCLE version 3.8.31 (Edgar, 2004), and trimmed using trimAl version v1.4.rev22 (Capella-Gutiérrez, Silla-Martínez and Gabaldón, 2009) to remove columns containing ⩾90% of gaps. The phylogeny was constructed using IQ-TREE multicore version 2.4.0 (Minh *et al.*, 2020) with the parameters of "-bb 1000 -m LG+G4". The phylogenetic tree was visualized in iTOL v7 (Letunic and Bork, 2019). The TerL fragments of Jug phages were concatenated and clustered using CD-HIT version 4.8.1(Huang *et al.*, 2010) with 99% identity, and only the representative sequences were included; the numbers in the brackets indicate the count of TerLs in the corresponding clusters.

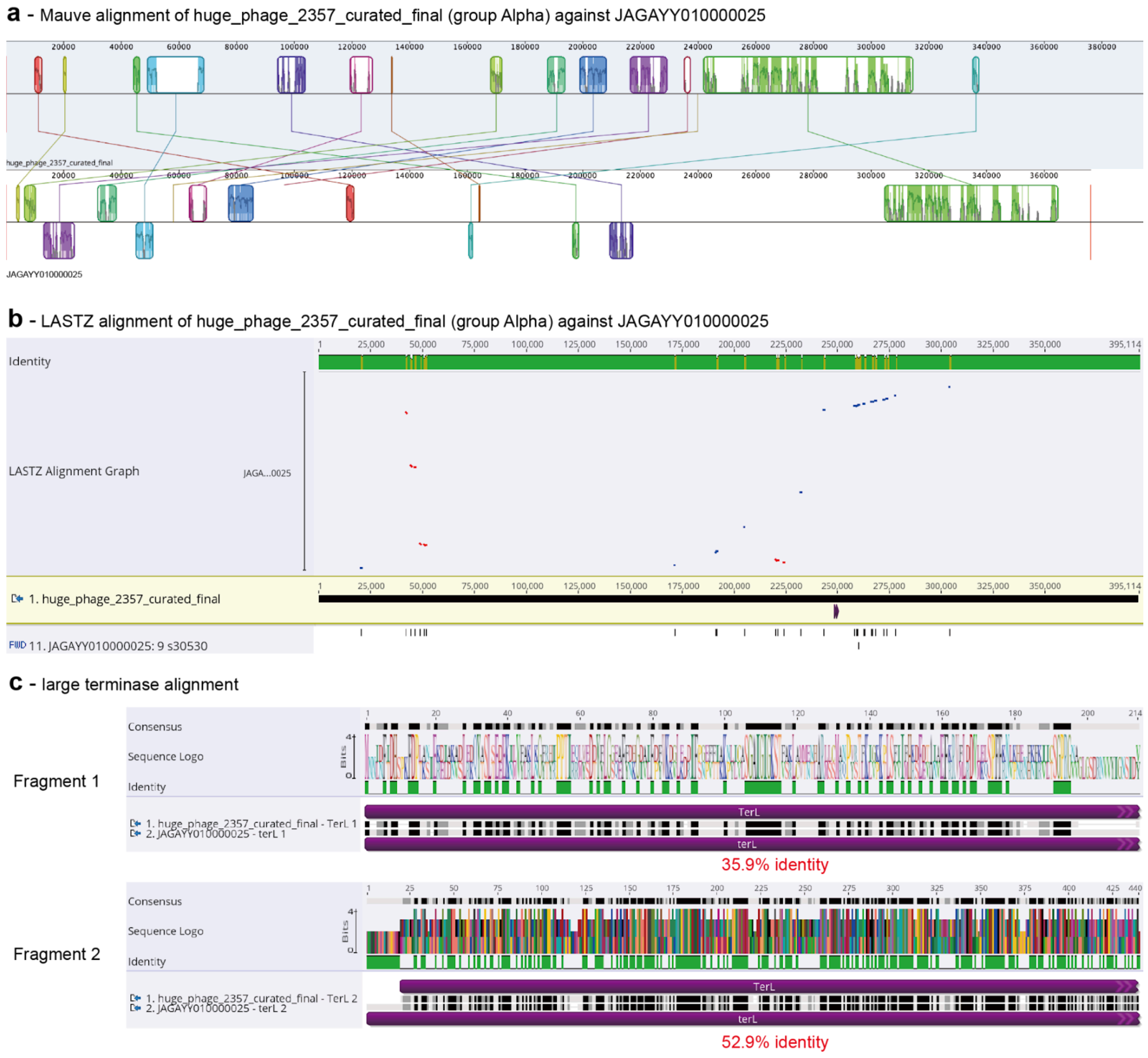


**Supplementary Figure 9 | The comparison of a phage genome (NCBI id = JAGAYY010000025) from the gut of water deer against the Jug phage.** When performed online BLASTp searches of Jug phage protein against NCBI-nr, we found that many of the Jug phage proteins had >50% identity to those from a metagenome-assembled genome (MAG), whose name is "Bacilli bacterium isolate RGIG8901 Water_deer_Omasum.Co__c192626, whole genome shotgun sequence". All the targeted hits were from a single contig in this MAG, JAGAYY010000025 (376,162 bp in length). We downloaded the sequence of JAGAYY010000025 and evaluated it using geNomad version 1.5.2 (Camargo *et al.*, 2023) and found that it represented a circular phage genome (with an end overlap sequence of 141 bp in length). This may indicate a misbinning of the phage contig into the MAG. The protein-coding genes of JAGAYY010000025 were predicted using Prodigal version 2.6.3 (Hyatt *et al.*, 2010) using the "single" mode, and searched against the proteins of the 139 curated Jug phages. Two large terminase (TerL) fragments were detected for JAGAYY010000025, and the longer one is most similar to that of huge_phage_2357_curated_final (group Alpha). We thus compared the genomes of huge_phage_2357_curated_final and JAGAYY010000025. (a) The Mauve alignment of the two genomes. It was analyzed using Mauve (Darling *et al.*, 2004). (b) The LASTZ alignment of the two genomes. It is analyzed using LASTZ version 1.04.15 (Harris, 2007). (c) The TerL alignments, which were conducted using MUSCLE version 5.1. All the figures were generated within Geneious Prime Build 2025-03-24 (Kearse *et al.*, 2012).

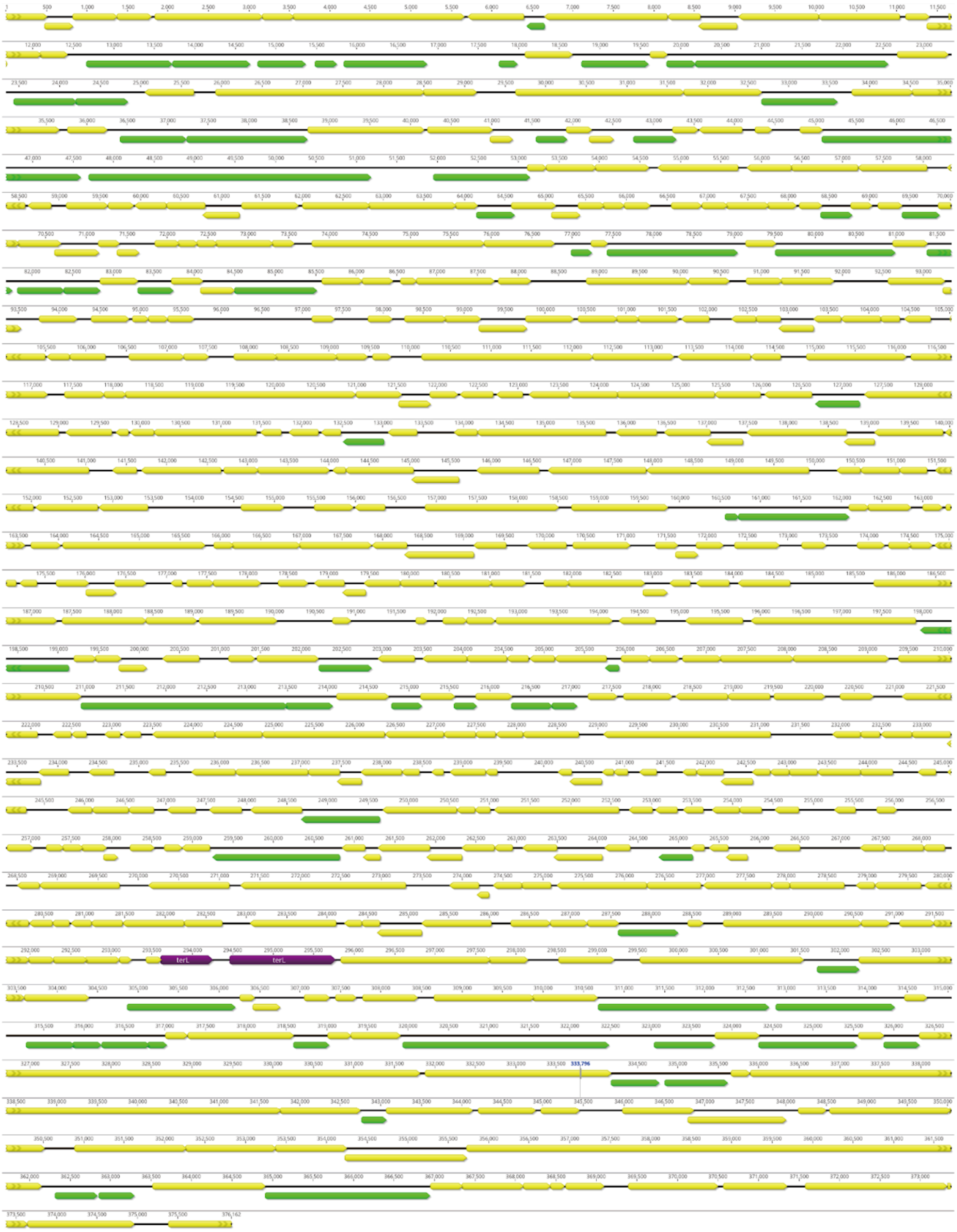


**Supplementary Figure 10 | The protein-coding genes of JAGAYY010000025.** The protein-coding genes shared >50% identity with those from any curated Jug phages are shown in purple (the two large terminase fragments) or green, with all other genes shown in yellow. The comparison of JAGAYY010000025 against the Jug phage protein-coding genes was performed using local BLASTp. The figure was generated within Geneious Prime Build 2025-03-24 (Kearse *et al.*, 2012).

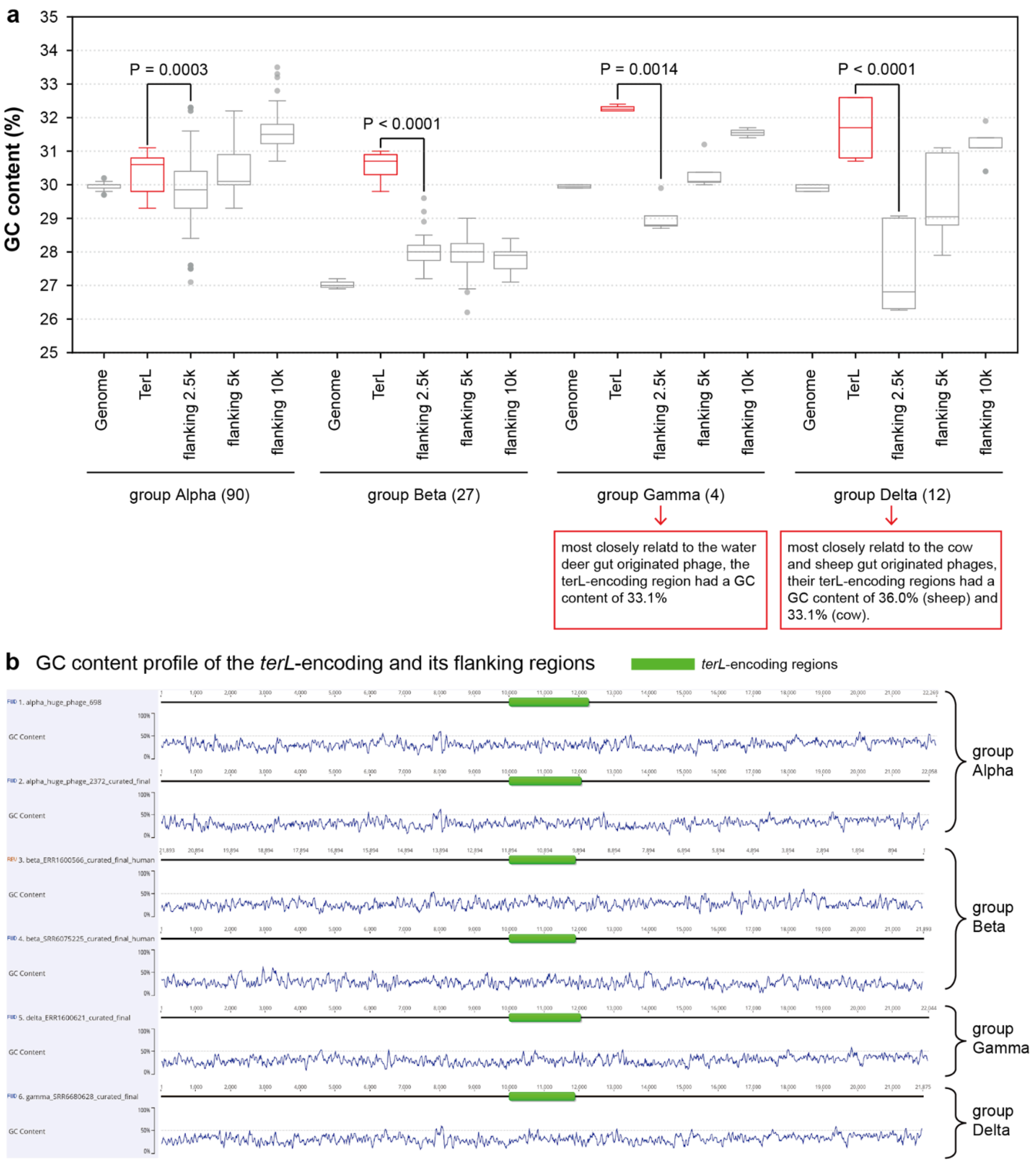


**Supplementary Figure 11 | The GC contents of the whole genome, TerL-encoding, and TerL-flanking regions.** (a) The comparison of GC contents. The flanking regions were extracted from the genomes based on the starting and ending positions of all the *terL* fragments. The significant difference in GC contents between the *terL*-encoding region and the flanking 2.5 kbp region was tested by a paired t-test. The flanking region, for example, 2.5 kbp, is defined as the concatenated nucleotide sequence of the upstream 2.5 kbp and downstream 2.5 kbp of the *terL*-encoding region, thus 5 kbp in total. (b) The examples of genomes from all four Jug phage groups show the GC content of the *terL*-encoding and its flanking regions.

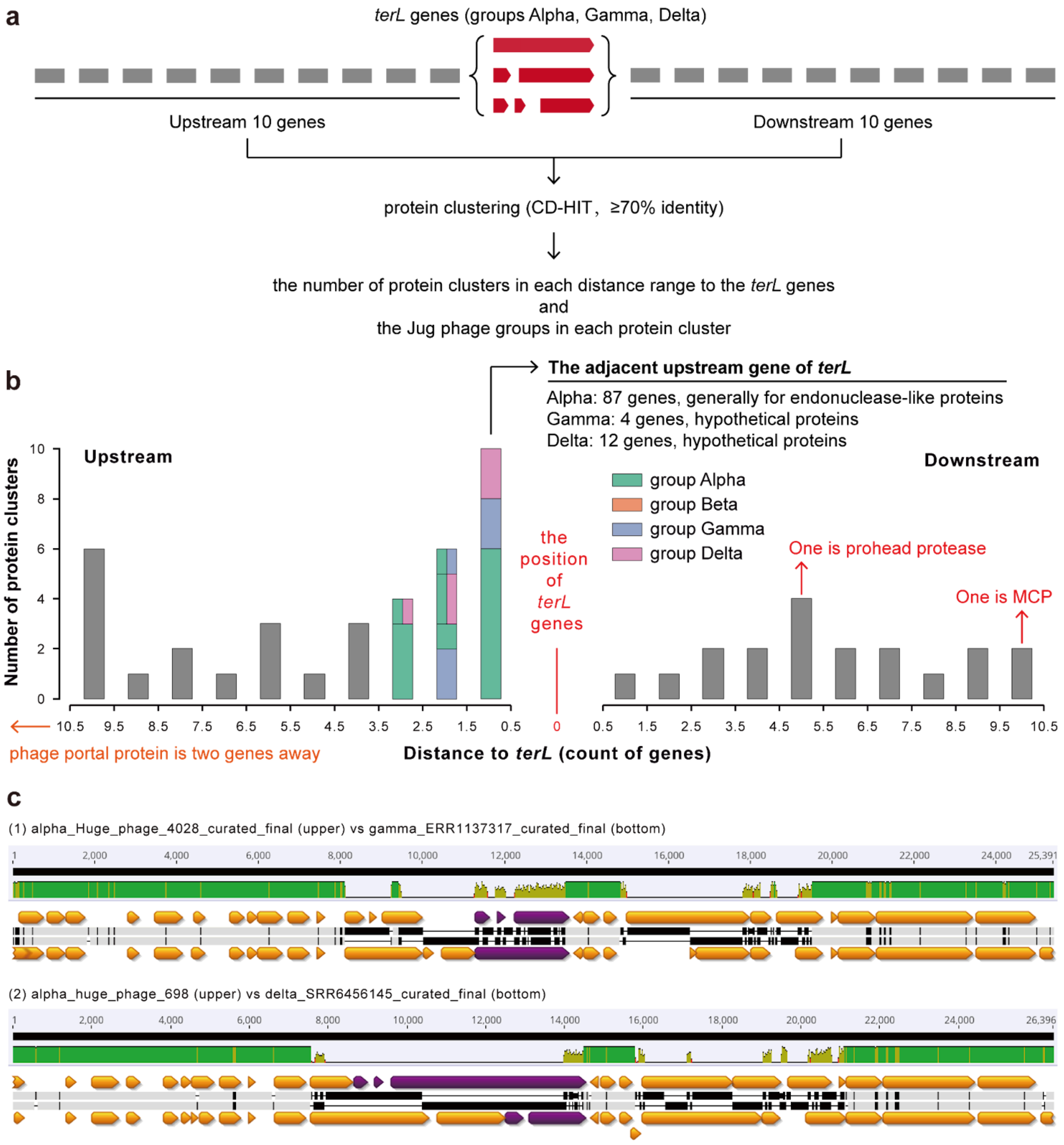


**Supplementary Figure 12 | The neighbor genes of the large terminase (*terL*) genes.** (a) The diagram shows the analysis pipeline of the neighborhood genes. The CD-HIT version 4.8.1 [66] was used to cluster the genes, with ⩾70% identity (-c 0.7 -aS 0.9 -G 0). Then, in each of the distance ranges, the number of protein clusters was summarized and presented in (b). (b) The number of protein clusters in each distance range to the *terL* genes. As we found that the protein clusters in the upstream distance range of 0.5-3.5 genes were very distinct, we thus profiled the corresponding Jug phage groups for each cluster. Notably, the upstream distance range of 0.5-1.5 genes contained 10 protein clusters, and those of group Alpha members generally encoded for endonuclease-like proteins, while those of groups Gamma and Delta were for hypothetical proteins. One of the protein clusters is for the major capsid protein (MCP). (c) The nucleotide alignment of the *terL*-coding regions of Jug phage genomes from different groups. Examples of (1) groups Alpha and Gamma, and (2) groups Alpha and Delta are shown. The TerL fragments are indicated in purple.

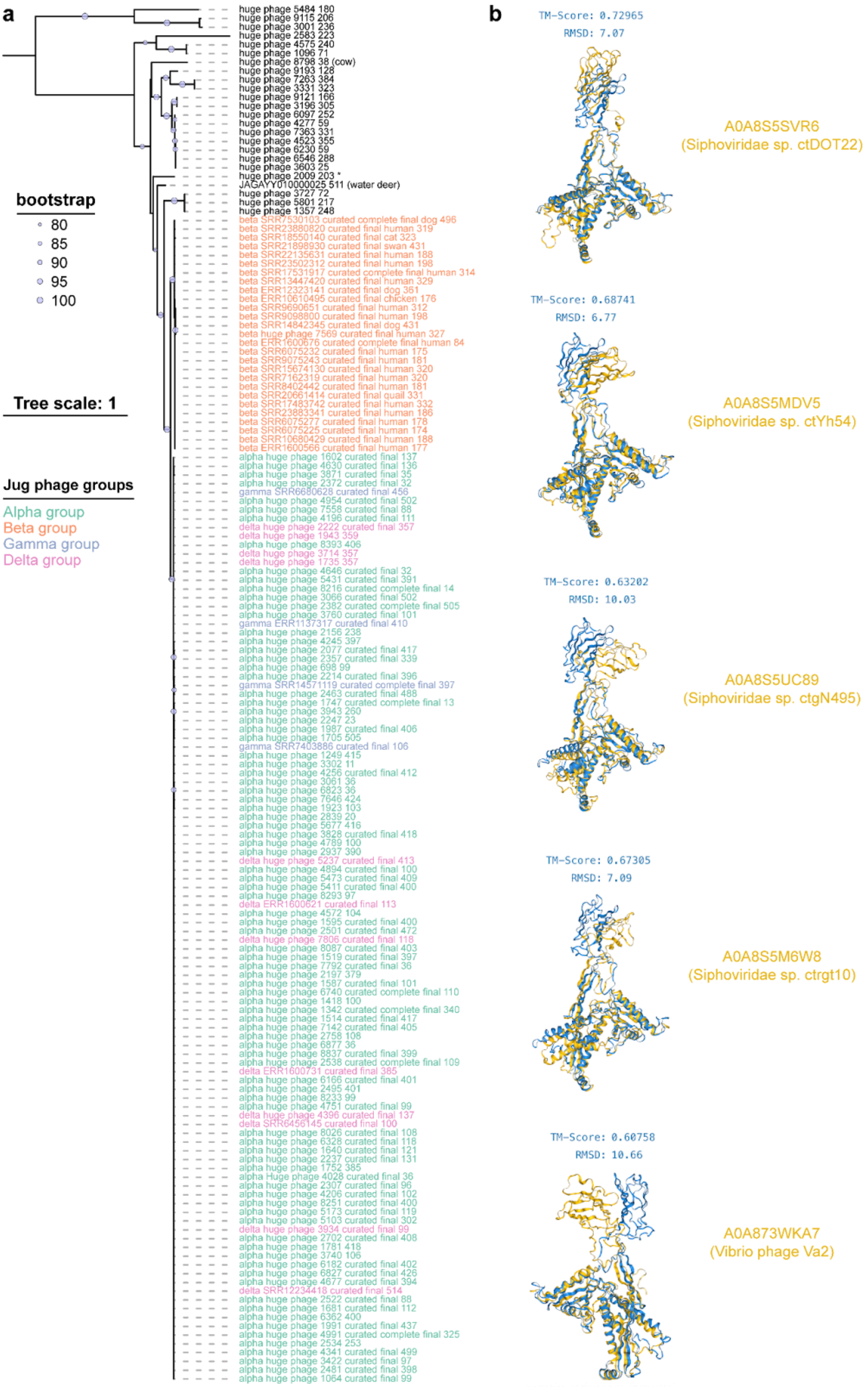


**See the text legend on the next page**

**Supplementary Figure 13 | The major capsid proteins of Jug phages.** (a) The phylogeny of the major capsid proteins. All the major capsid proteins (MCPs) from Jug phages and their relatives were included for analysis. The MCPs encoded by relatives were retrieved by searching all other HPGC proteins against the Jug phages' MCPs using BLASTp with an e-value threshold of 1e-5. The MCP sequence of the phage detected in the water deer gut microbiome was included as well. All the Jug phage relatives were from the gut microbiomes, except for the one from a wastewater sample (indicated by *). (b) The comparison of the MCP structures. The top five BFVD MCP structure hits with the highest TM-scores against Jug phages' MCP are shown. The BFVD/UniProt IDs of the reference structures and the corresponding viruses are shown on the right.

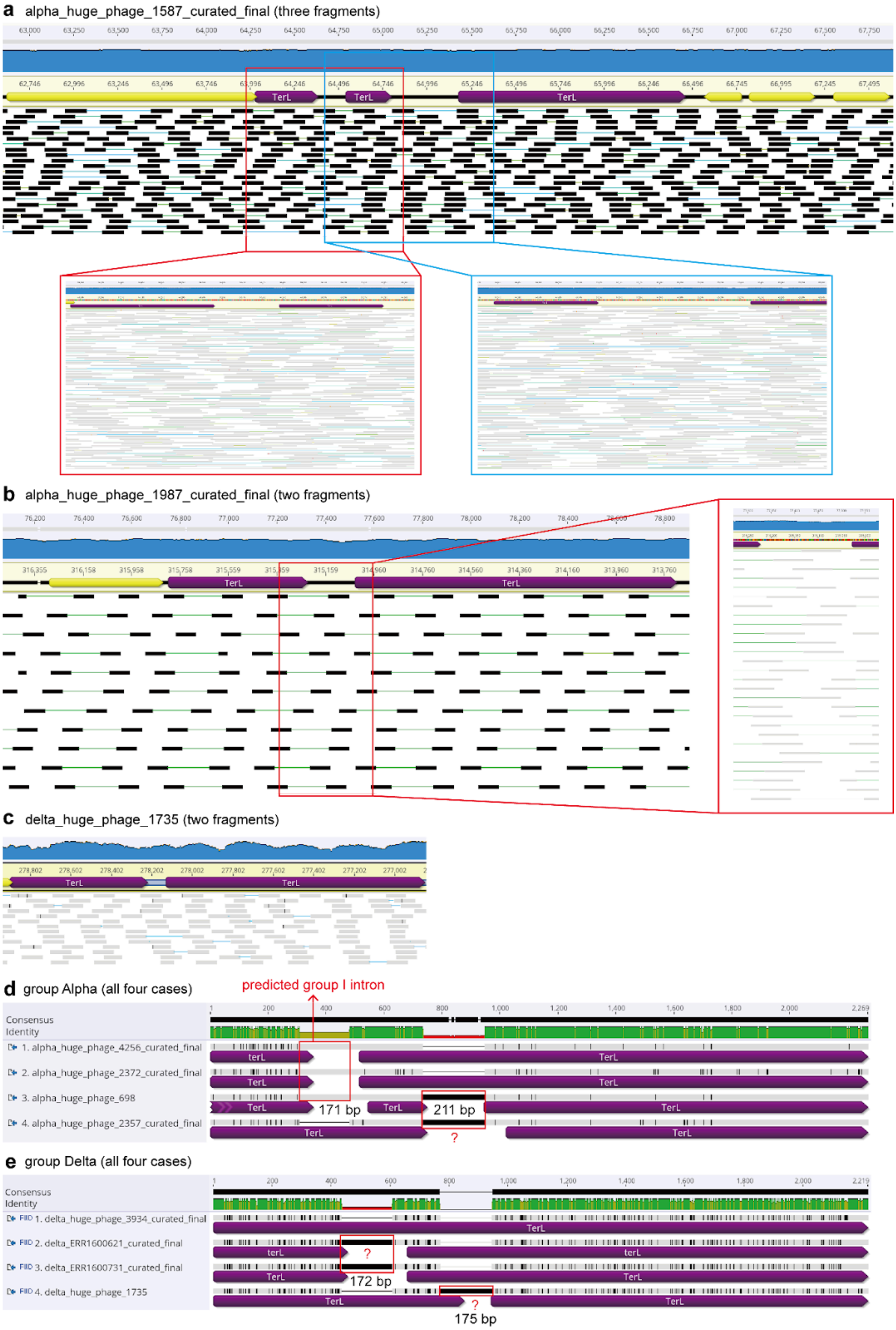


**See the text legend on the next page**

**Supplementary Figure 14 | The fragmentation of the large terminase (*TerL*) genes encoded by Jug phages.** In (a), (b), and (c), examples of paired-end reads mapping to the *TerL*-encoding regions did not show any single-nucleotide indel that led to the fragmentation of the *TerLs*. The zoom-in of the mapped paired-end reads is shown in the red and/or blue boxes. In (d) and (e), alignment of the nucleotide sequences of the *TerL*-encoding regions likely indicated that the insertion of large nucleotide sequences could be the potential reason for *TerL* fragmentation. The nucleotide length of the introns is shown. The mapped was performed using Bowtie2 version 2.4.4 (Longmead and Salzberg, 2012) with default parameters, and the figures were generated using Geneious Prime Build 2025-03-24 (Kearse *et al.*, 2012).

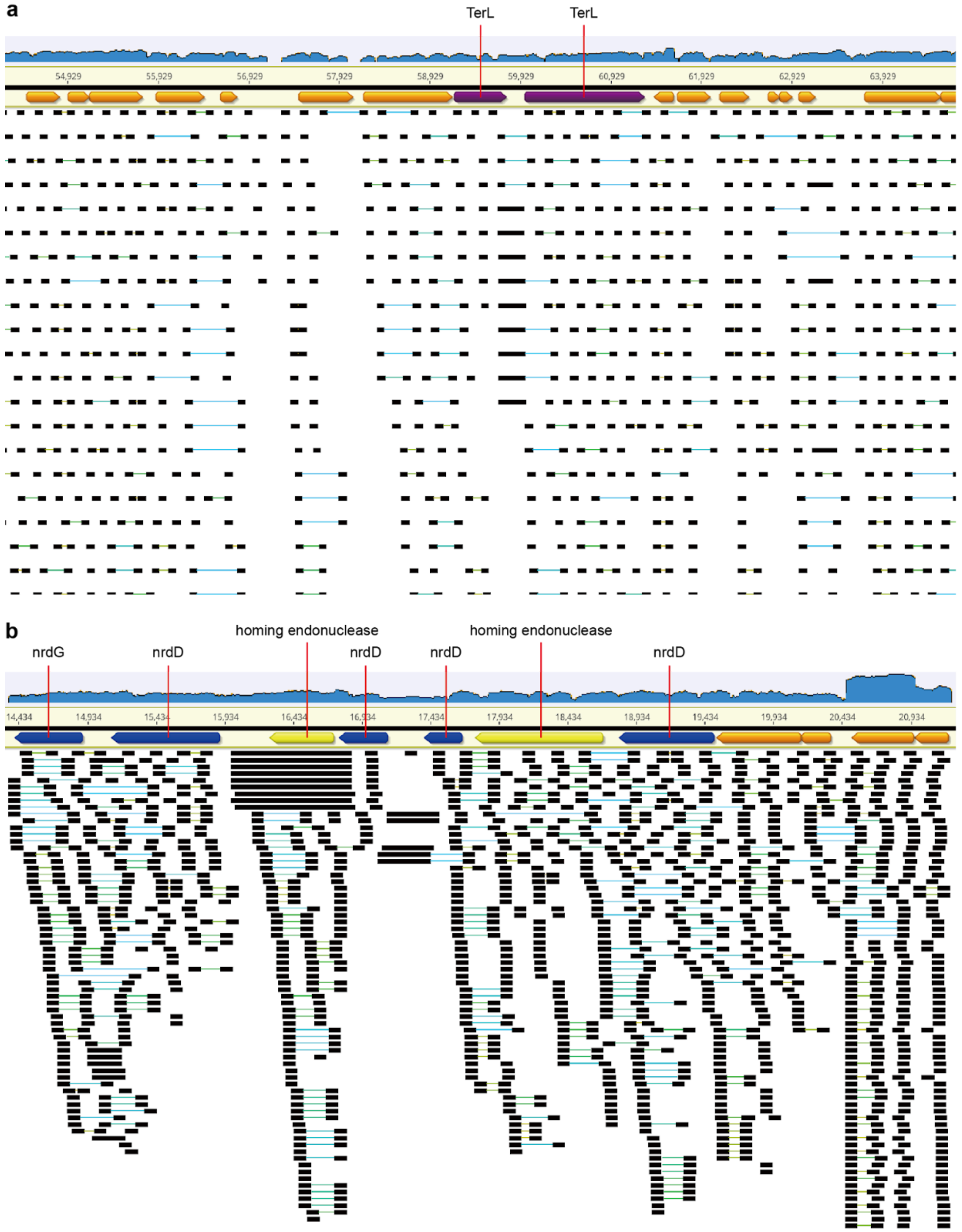


**Supplementary Figure 15 | The co-transcription of fragmented genes as operons, with (a) the large terminase (TerL) and (b) the nrdD as examples.** The RNA reads from the adult male 1 samples were mapped to the corresponding Jug phage genome. The RNA reads mapping bam file was imported and visualized using Geneious Prime Build 2025-03-24 (Kearse *et al.*, 2012).

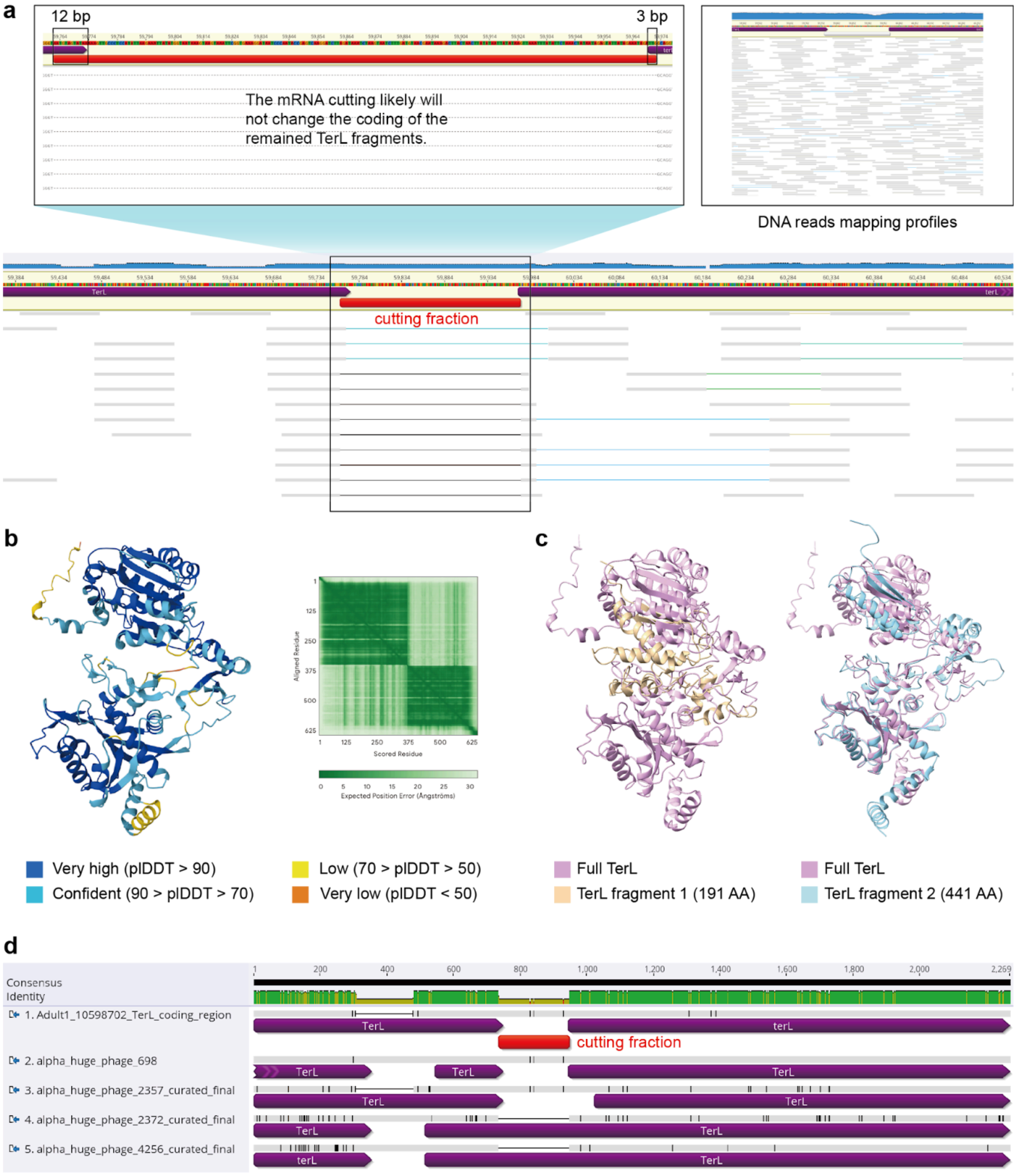


**Supplementary Figure 16 | An example showing the mRNA cutting and splicing to obtain a full-length TerL.** The terL fragments of Jug phage from adult 1 are used as an example here. (a) The mRNA cutting won't change the coding of the remained fragments. The DNA reads mapping profiles are shown on the right, to indicate that the RNA mapping pattern was not due to a local variation in which some phage genomes lacked the "cutting fraction" region. (b) The TerL protein structure predicted by AlphaFold3 (https://alphafoldserver.com/). With the pLDDT values of each residue shown on the left, and the alignment information on the right. (c) The structure alignment of the TerL fragments to the full-length TerL. (d) The nucleotide alignment of the terL-coding regions of curated Jug phage group Alpha genomes with that of adult 1. The alignment supports our conclusion of mRNA cutting and splicing based on RNA reads mapping. See **Supplementary Figure 14** for more information.

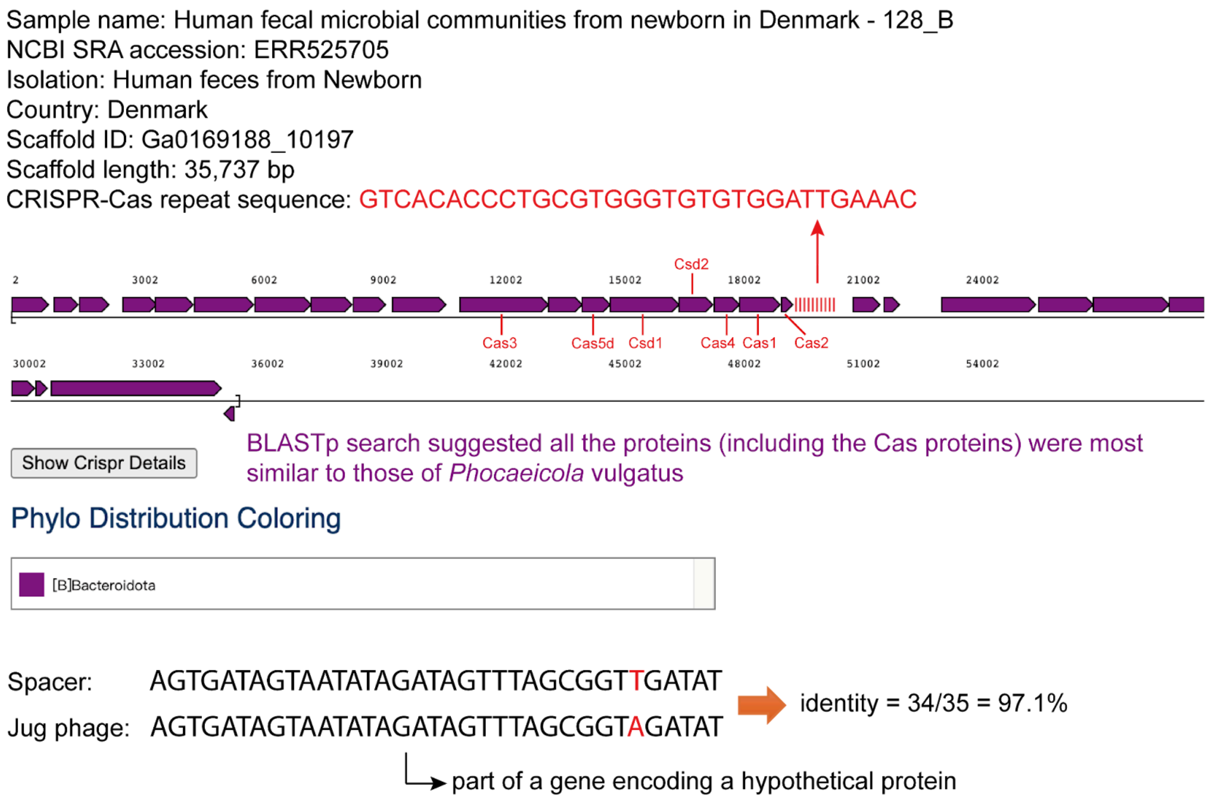


**Supplementary Figure 17 | An example of CRISPR-Cas spacer targeting on the Jug phage genomes.** The information about the scaffold with the CRISPR-Cas system is shown. The scaffold was assembled from the metagenomic reads of a newborn gut microbiome sample, and its taxonomic assignment is *Phocaeicola vulgatus*. The diagram in the middle shows the locations of the cas proteins and the repeat locus on the scaffold. The alignment of the targeting spacer and the Jug phage genomic fragment is shown at the bottom.

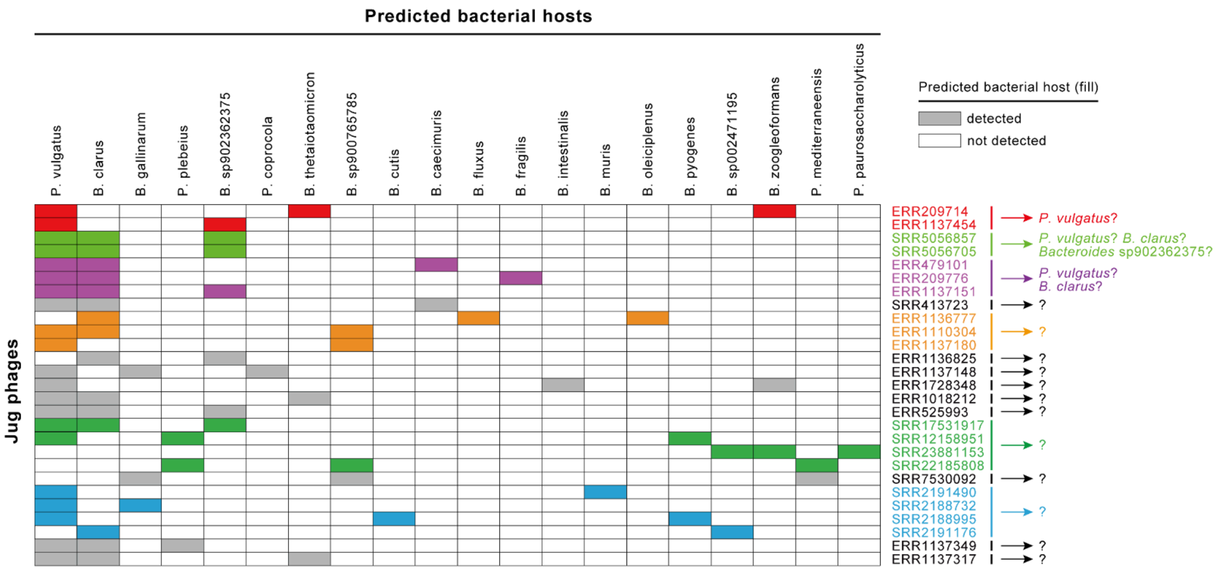


**Supplementary Figure 18 | The co-occurrence of *Bacteroides* and *Phocaeicola* species and the Jug phages.** Only those samples with only 2 or 3 predicted hosts (*Bacteroides* and *Phocaeicola* species) detected are shown. Each line represents a Jug phage (indicated by the sample SRA numbers, as only one Jug phage is presented in the corresponding samples), and each column represents a *Bacteroides* or *Phocaeicola* species. The SRA numbers in the same color (excepting black) indicated that the Jug phage genomes shared ⩾97% genome similarity across ⩾90% of their genome length (by dRep version 3.2.2 (Olm *et al.*, 2017)), and such phages were assumed to infect the same host(s). The colored boxes indicate the presence of the *Bacteroides* or *Phocaeicola* species in the corresponding samples. The specific host-virus relationship was speculated based on co-occurrence and is shown on the right.

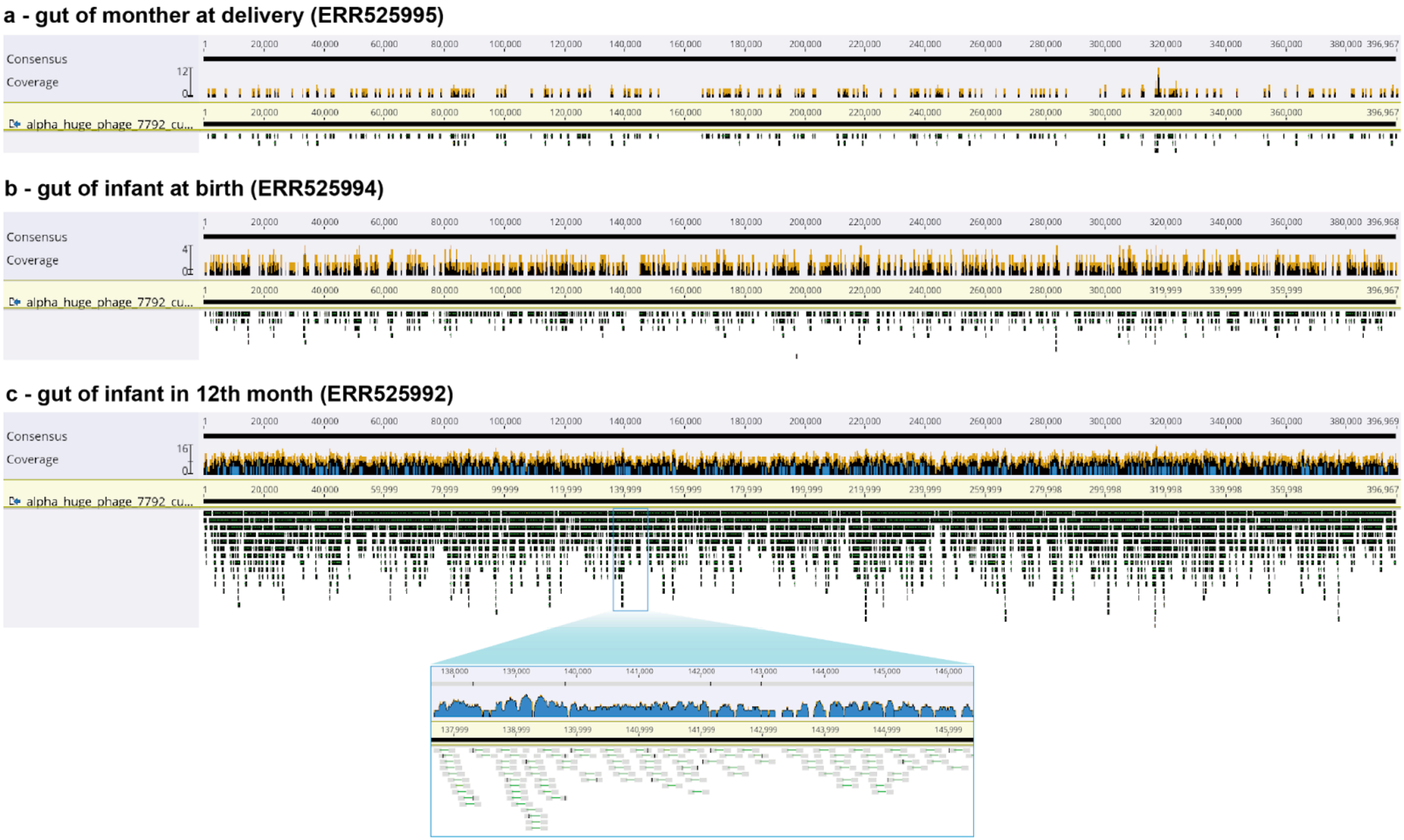


See text legend on the next page.

**Supplementary Figure 19 | The detection of the group Alpha Jug phage in the gut microbiomes of the infant and the mother.** The mapping of reads from gut samples collected (a) at delivery of the mother, (b) at birth of the infant, and (c) in the 12th month of the infant. The paired-end reads were mapped to the curated Jug phage genome reconstructed from the gut samples collection in the fourth month of the infant, allowing no more than one mismatch for each read to be mapped. The mapping was performed using Bowtie2 version 2.4.4 (Longmead and Salzberg, 2012) with default parameters, and the figures were generated using Geneious Prime Build 2025-03-24 (Kearse *et al.*, 2012). The zoom-in indicates that each black square represents a metagenomic read. The paired-end reads are linked with a blue line.

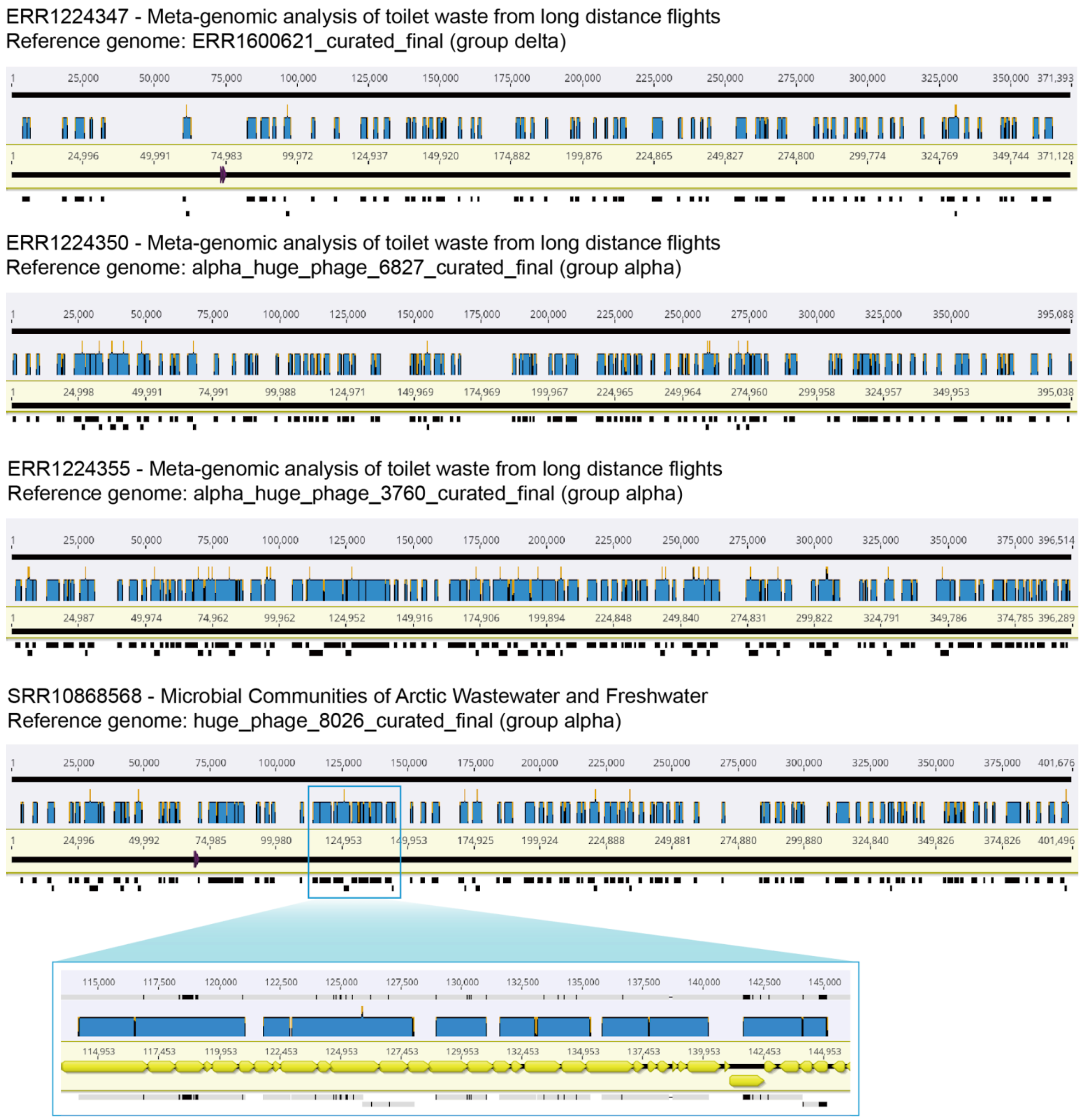


**Supplementary Figure 20 | Genome alignment of Logan contigs assembled from wastewater to the curated Jug phage genomes.** The alignment was performed using the "Map to Reference(s)" function of Geneious Prime Build 2025-03-24 (Kearse *et al.*, 2012). Four examples (one of group Delta, and three of group Alpha) are shown with the sample information, including SRA IDs and project descriptions from NCBI. The zoom-in shows the details of the alignment as an example.

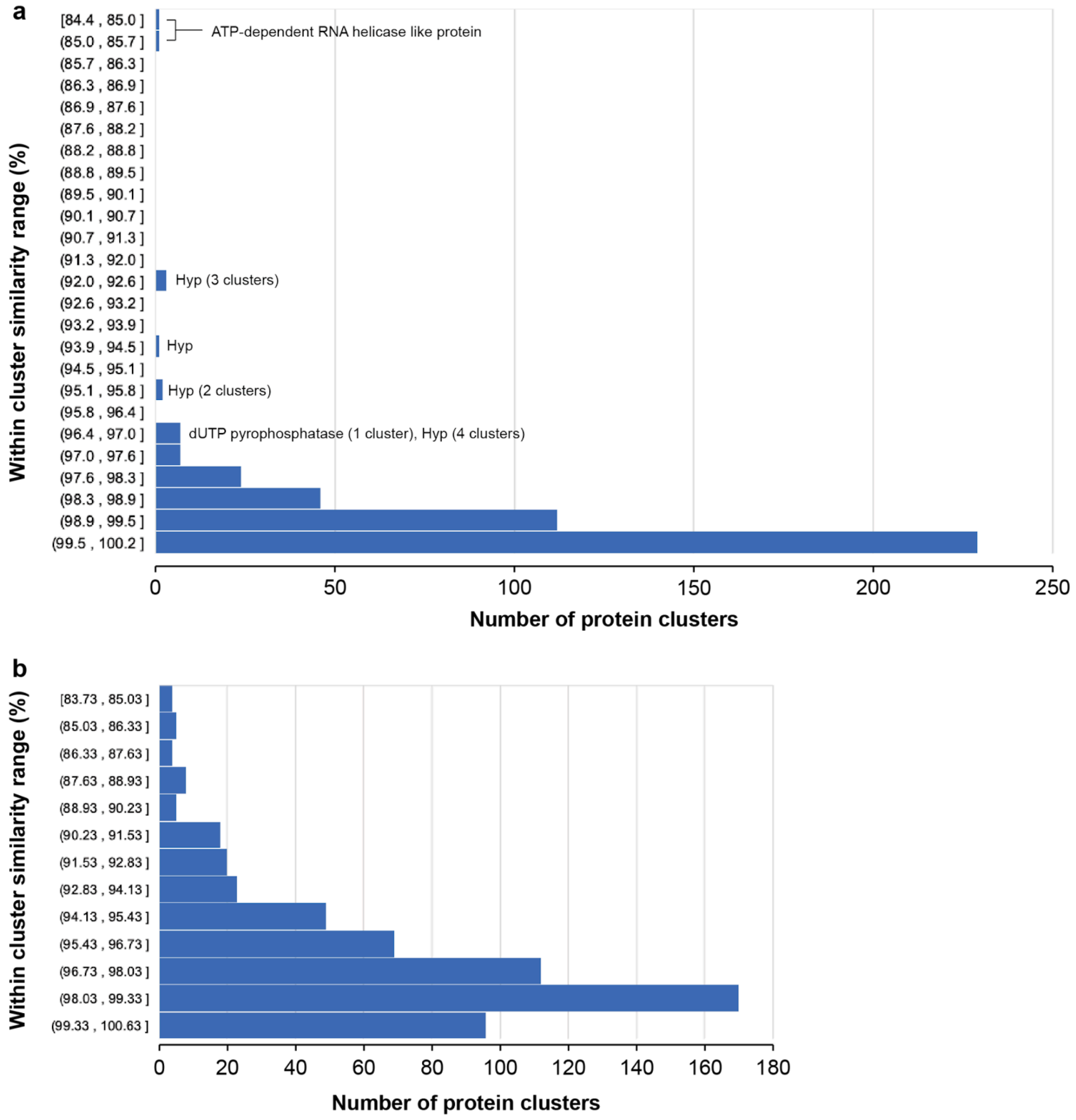


**Supplementary Figure 21 | The similarity of proteins within each protein cluster of the Jug phages.** The protein clusters shared by (a) the group Alpha Jug phages from the gut of humans, mice, and rats, and (b) the group Beta Jug phages from the gut of humans and dogs. The average paired protein sequence identity of each cluster was calculated first, and then the similarity ranges of all shared protein clusters were profiled. The paired protein sequence similarity was based on all-vs-all BLASTp within each cluster (e-value threshold = 1e-5).

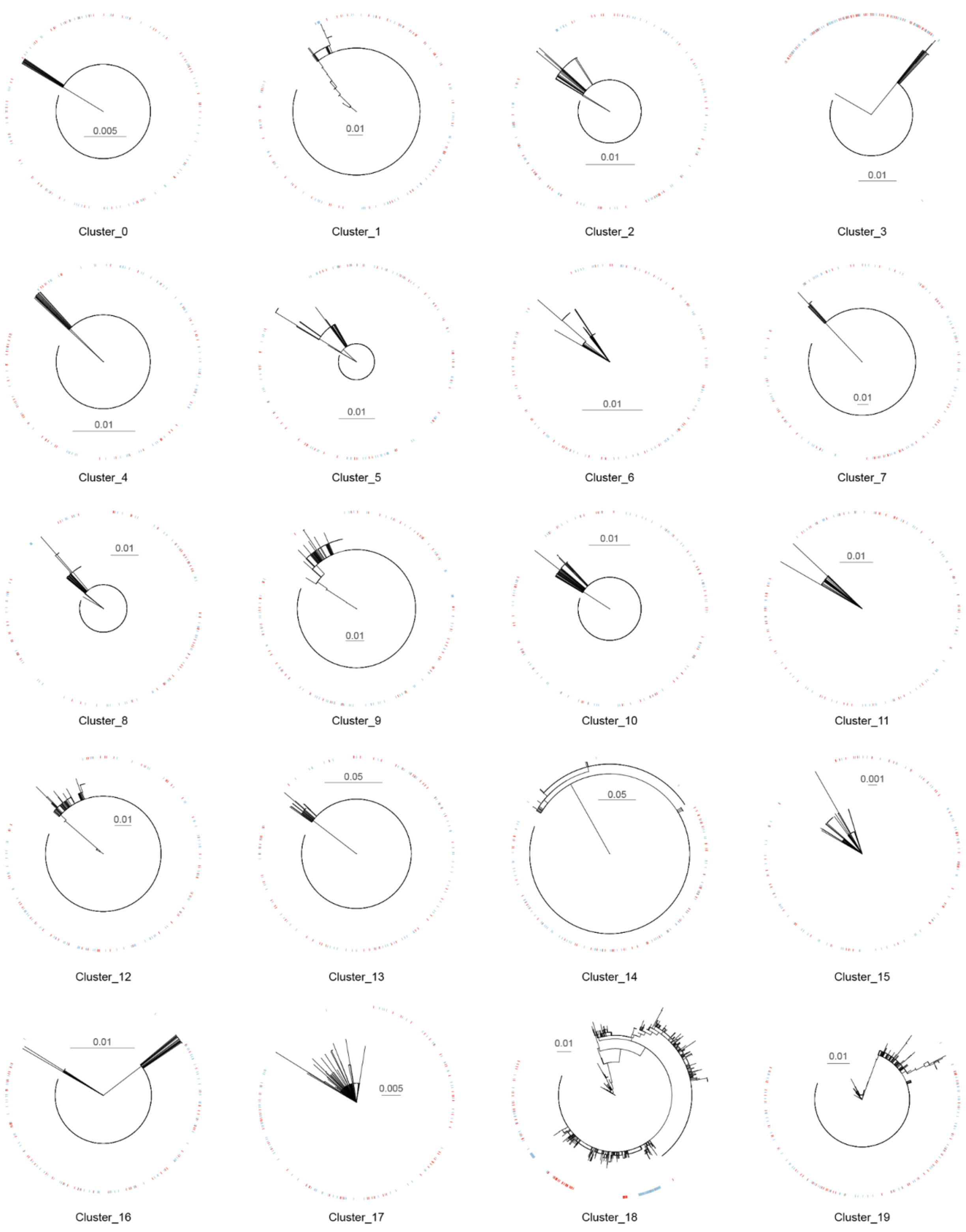


**Supplementary Figure 22 | The phylogenies of protein families shared by group Apha Jug phages from humans, mice, and rats.** The sequences of the protein families containing group Alpha Jug phages from the gut of humans, mice, and rats were conducted for phylogenetic analyses, and the top 20 protein clusters are shown here as examples. The colored strips indicate the animal host of the Jug phages, with blue strips for those from mice, red strips for those from rats, and all others for those from humans.

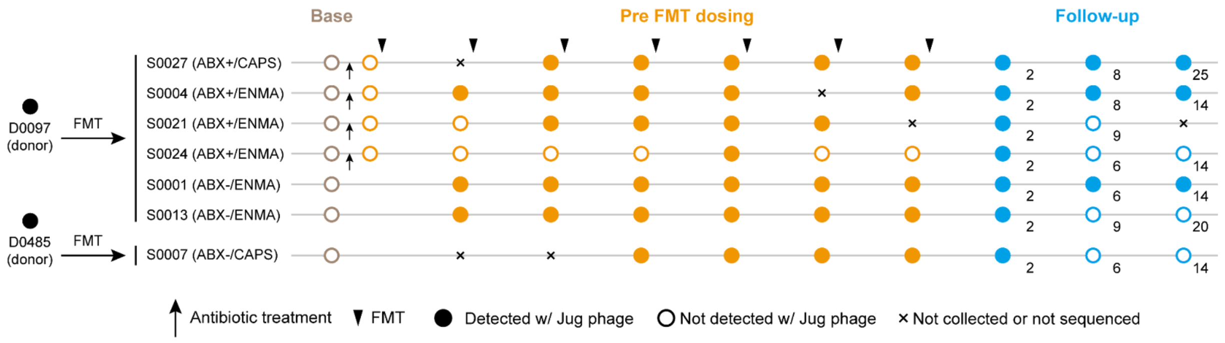


**Supplementary Figure 23 | The fecal microbiota transplantation (FMT) from two donors to the recipients for the treatment of ulcerative colitis.** The points of recipients that received antibiotic treatment before FMT are indicated by black arrows. A total of 7 FMT dosing and 2-3 follow-up evaluations were performed for each recipient. The colored circles indicated when the fecal samples were collected for metagenomic analyses; one more sample was collected for those recipients accepting antibiotic treatment. The follow-up time (in weeks) after the last dosing is shown. The FMT was performed in two approaches, i.e., ENMA (maintenance doses via enema) and CAPS (maintenance doses via capsules). ABX+, with antibiotic pretreatment; ABX-, without antibiotic pretreatment.

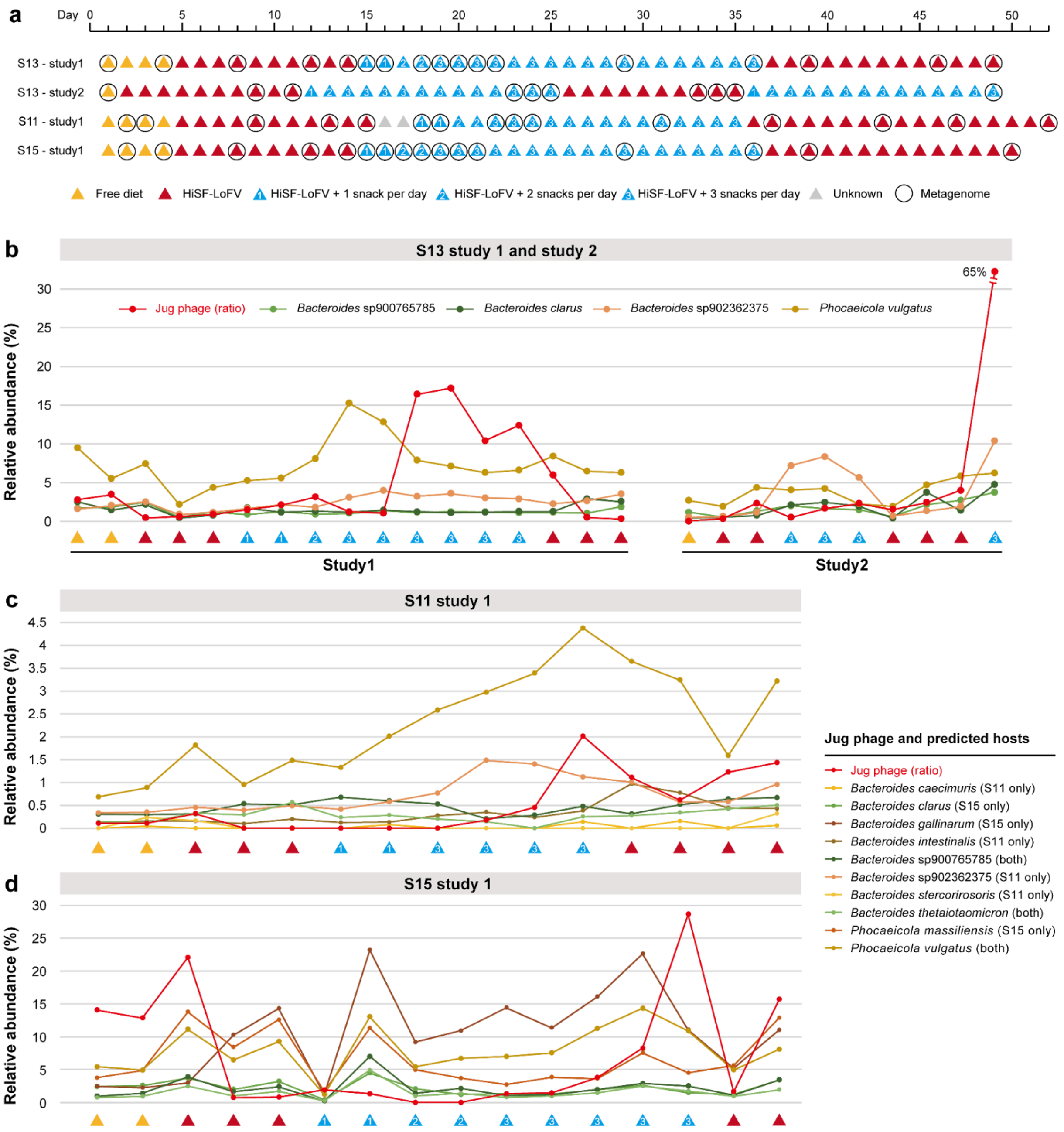


**Supplementary Figure 24 | The dynamics of the Jug phage in the three participants involved in the dietary intervention studies.** (a) The set-up of the dietary intervention study. The diet contents and metagenomic sampling time points (black circles) are shown. In study 1, the participants took a pea fibre-containing snack as a supplement, and had a 2-fibre (from day 12 through 25) or a 4-fibre snack prototype (from day 36 through 49). (b)-(d) The co-occurrence of Jug phages and *Bacteroides* and *Phocaeicola* species in the human gut based on sequencing coverage. The bacterial relative abundance was calculated based on the sequencing depth of the ribosomal protein S3 (rpS3) encoding scaffolds (see **Methods** in the main text). In (c) and (d), the presence of the predicted hosts in the participants is indicated in the brackets; "both" means the species was presented in both participants. For comparative analysis, we also show the presence of Jug phage by calculating the ratio of Jug phage sequencing coverage to the cumulative sequencing coverage of all bacterial and archaeal rpS3-encoding scaffolds within each respective sample.

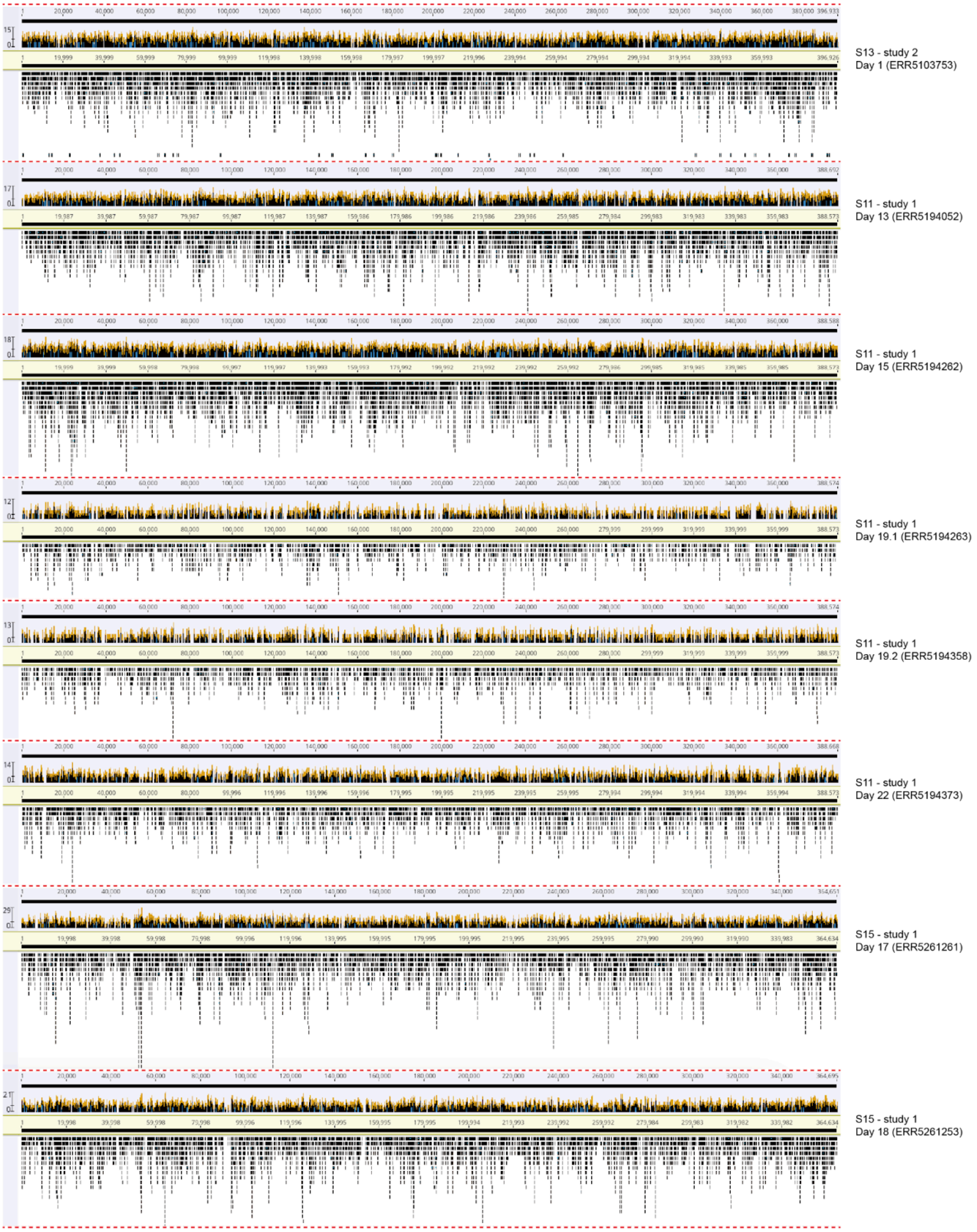


**Supplementary Figure 25 | The detection of the Jug phages in the gut microbiomes of the participants in dietary intervention studies.** The quality-control metagenomic paired-end reads were mapped to the corresponding curated Jug phage genomes reconstructed for each participant, allowing no more than one mismatch for each read to be mapped. The mapping was performed using Bowtie2 version 2.4.4 (Longmead and Salzberg, 2012) with default parameters, and the figures were generated using Geneious Prime Build 2025-03-24 (Kearse *et al.*, 2012).

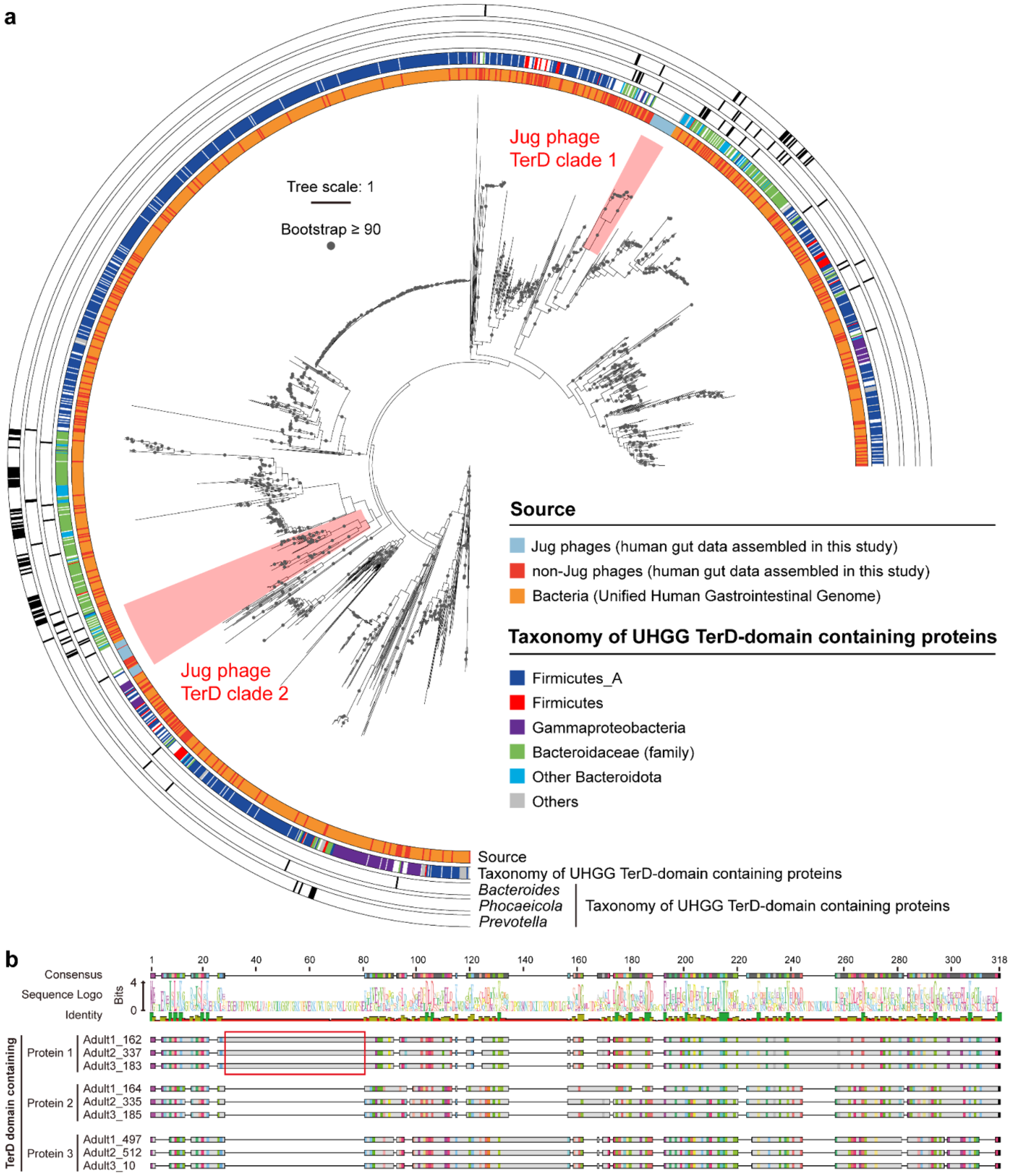


**Supplementary Figure 26 | The TerD domain-containing genes encoded by Jug phages.** (a) The phylogeny of the TerD domain-containing proteins. The source of the proteins (the most inner colored strip circle) includes (1) Jug phages, and (2) non-Jug phages with sequences assembled in this study, and the non-Jug phage ones from the Unified Human Gastrointestinal Genome (UHGG) unique protein database. The proteins from each source were firstly clustered with ⩾99% identity using CD-HIT, and the cluster representatives were included for phylogenetic analyses. The taxonomy of the UHGG TerD-domain containing proteins is shown in the second most inner colored strip circle; note that those of the family Bacteroidaceae (containing the genera of *Bacteroides*, *Phocaeicola*, and *Prevotella*) are shown separately. The outer three colored strip circles indicate the TerD domain-containing proteins from *Bacteroides*, *Phocaeicola*, and *Prevotella*, respectively. The two clades with TerD proteins from the Jug phages are highlighted with a colored (red) background. (b) The alignment of the TerD protein sequences. Each of the three Jug phages encoded three copies of the *terD* genes. One of the TerD proteins has an extra region (indicated by the red box).

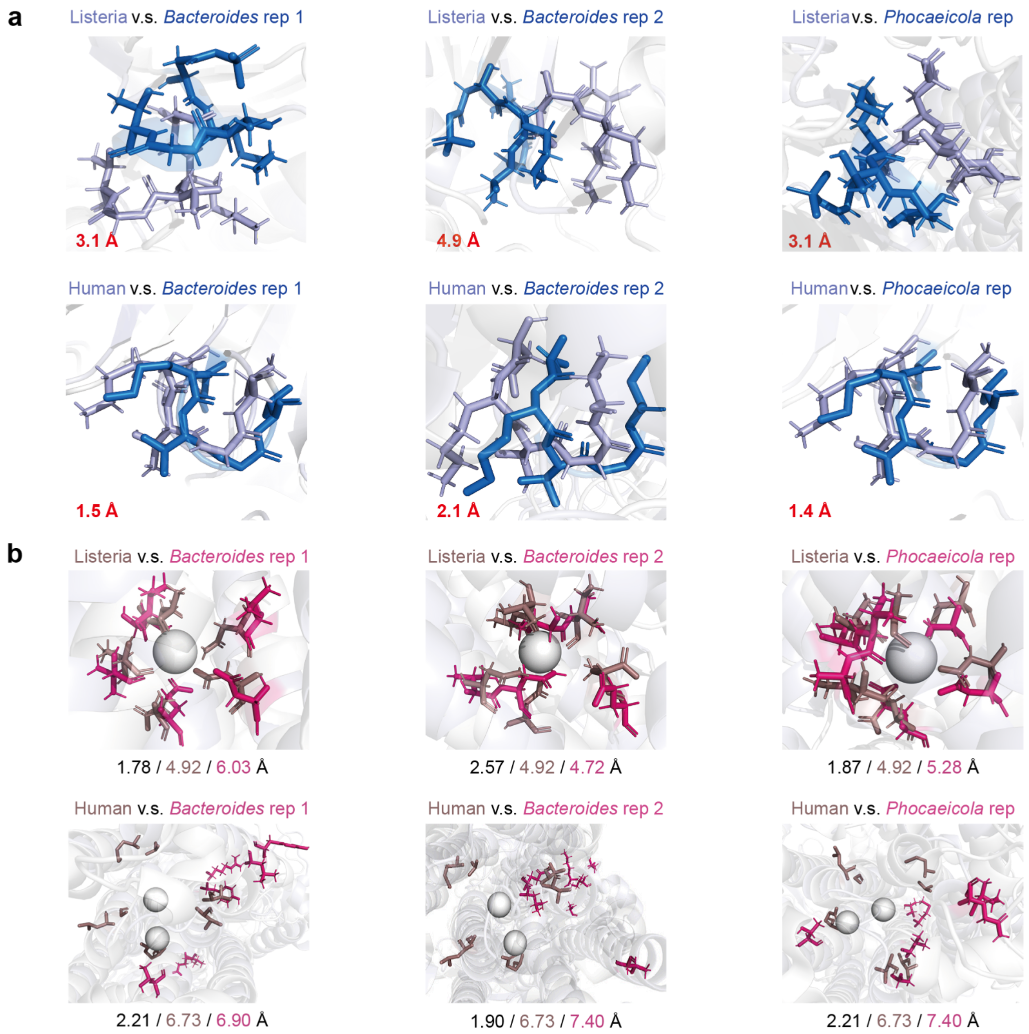


**Supplementary Figure 27 | The comparison of the calcium-translocating P-type ATPases encoded by predicted bacterial hosts of Jug phages against references.** (a) The comparison of the "DKTGT" motif. The distance (Å) of the "D" residue is shown in red. (b) Calcium coordination site comparison. The total distance of $Ca^{2+}$ ion (gray spheres) binding-related residues between the ATPases of predicted Jug phages' hosts and reference, the total distance of the reference ATPase binding residues to $Ca^{2+}$ ion, and the total distance of predicted hosts' ATPase binding residues to $Ca^{2+}$ ion are shown at the bottom. Note that the human ATPase could combine with two $Ca^{2+}$ ions. See Methods in the main text for more details.

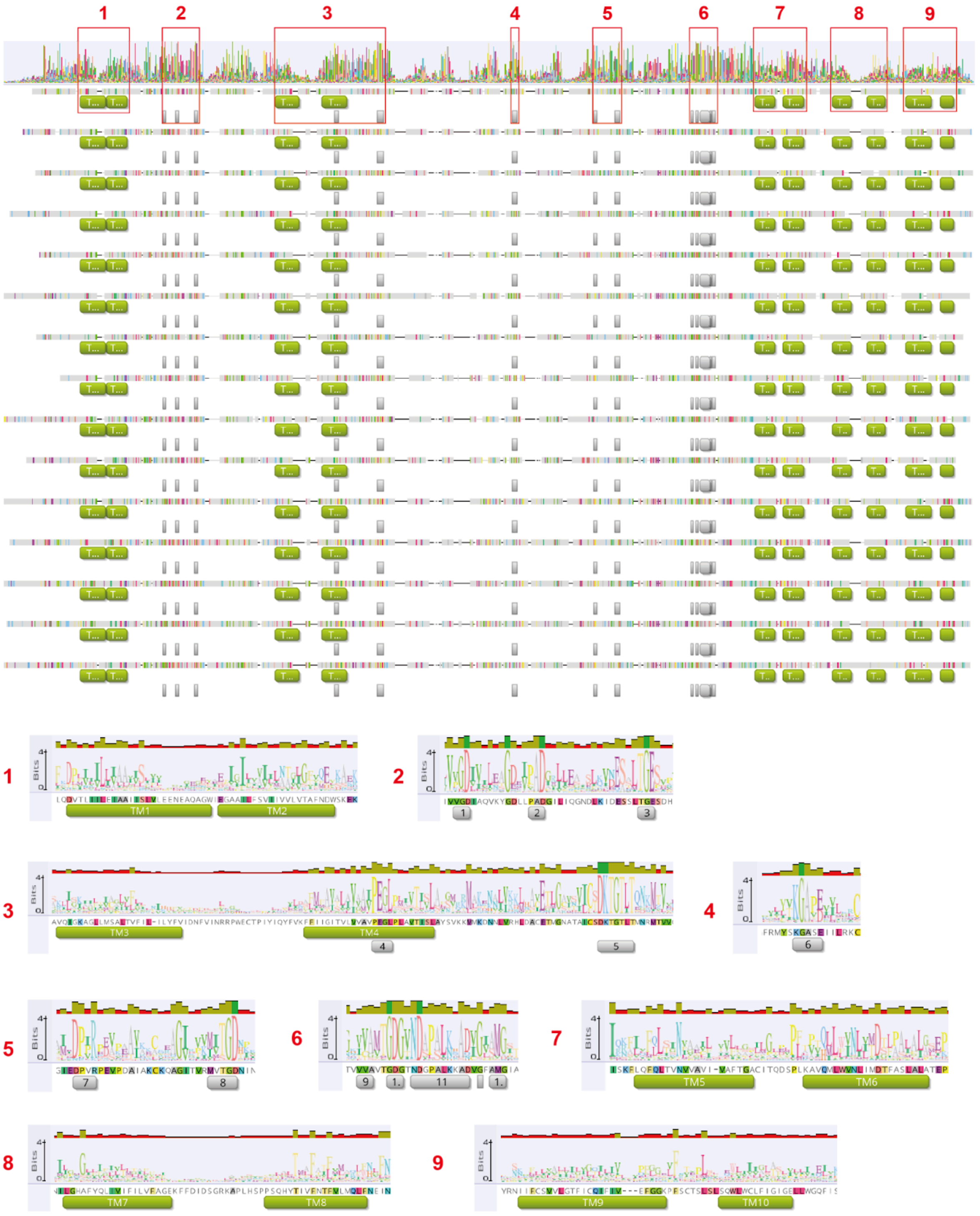


**Supplementary Figure 28 | The sequence alignment and key residues of calcium-tranlocating P-type ATPases.** The aligned sequences of the calcium-translocating p-type ATPases in **Supplementary Figure 28**, including those from *Homo sapiens* (ATP2B4), the *Chlorella* viruses (M535L and C785L), the *Listeria monocytogenes* (LMCA1), the Jug phages, the Jug phage relative from the water deer gut, and the co-existed gut bacteria of the three adult male were visualized using Geneious Prime Build 2025-03-24 (Kearse *et al.*, 2012). Only a subset of the aligned sequences is shown here. The locations of the 10 transmembrane regions and conserved residues were manually inspected and indicated based on the information from Bonza et al. (Bonza et al. 2010). These regions and conserved residues were zoomed in below, with the sequence logo shown.

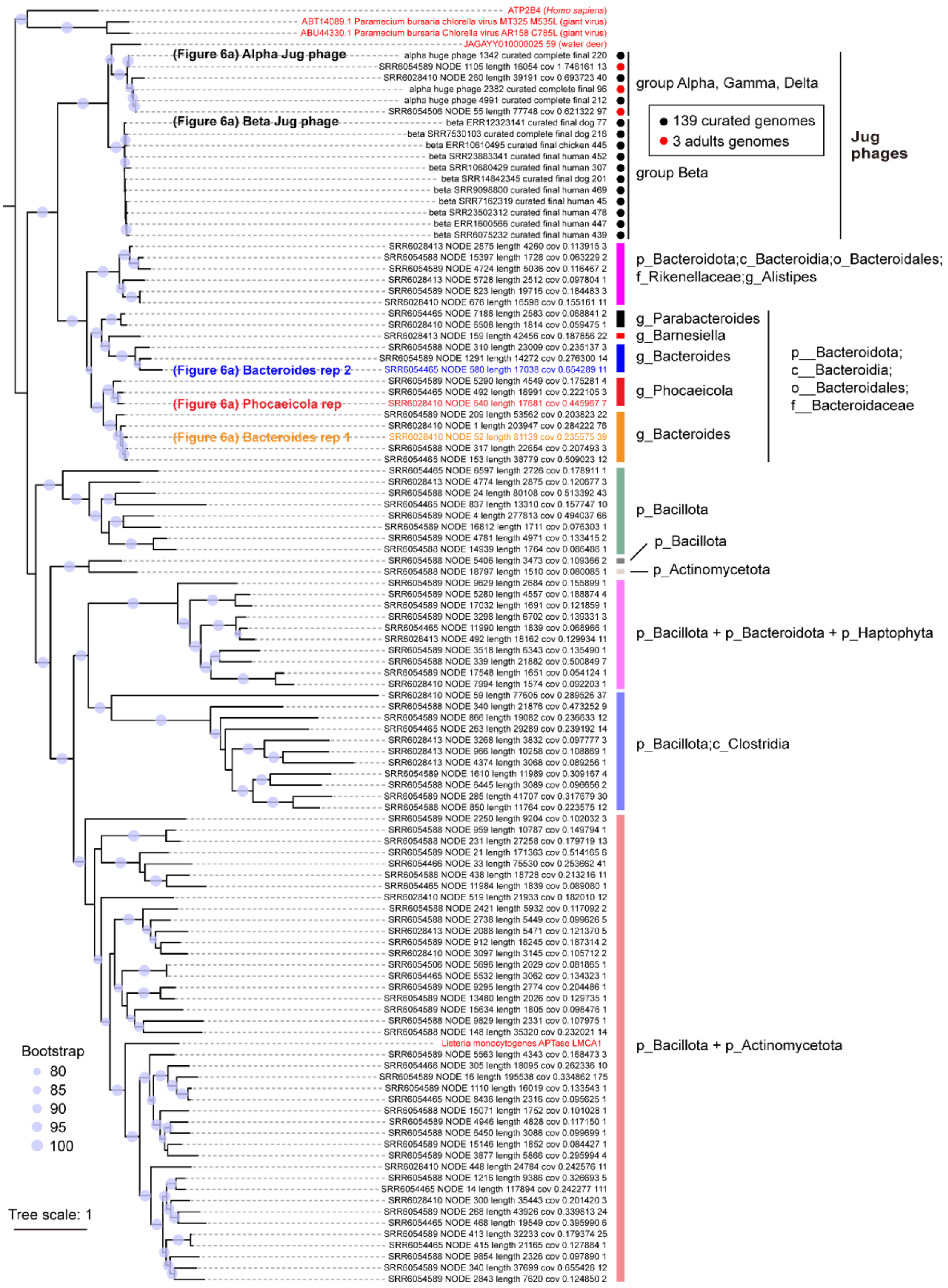


**Supplementary Figure 29 | The phylogeny of calcium-translocating P-type ATPases of Jug phages and human gut bacteria.** The calcium-translocating P-type ATPases included those from *Homo sapiens* (ATP2B4), the *Chlorella* viruses (M535L and C785L), the *Listeria monocytogenes* (LMCA1), the Jug phages and their relative from the water deer gut, and the co-existed gut bacteria of the three adult males. Those sequences from the curated Jug phage genomes were dereplicated using CD-HIT by clustering them with 99% identity, and only the cluster representatives were included. The taxonomy of scaffolds encoding the bacterial ATPases was determined based on the taxonomic assignment of all protein-coding genes on the corresponding scaffolds. The ones predicted with protein structures and shown in Figure 6a are indicated.

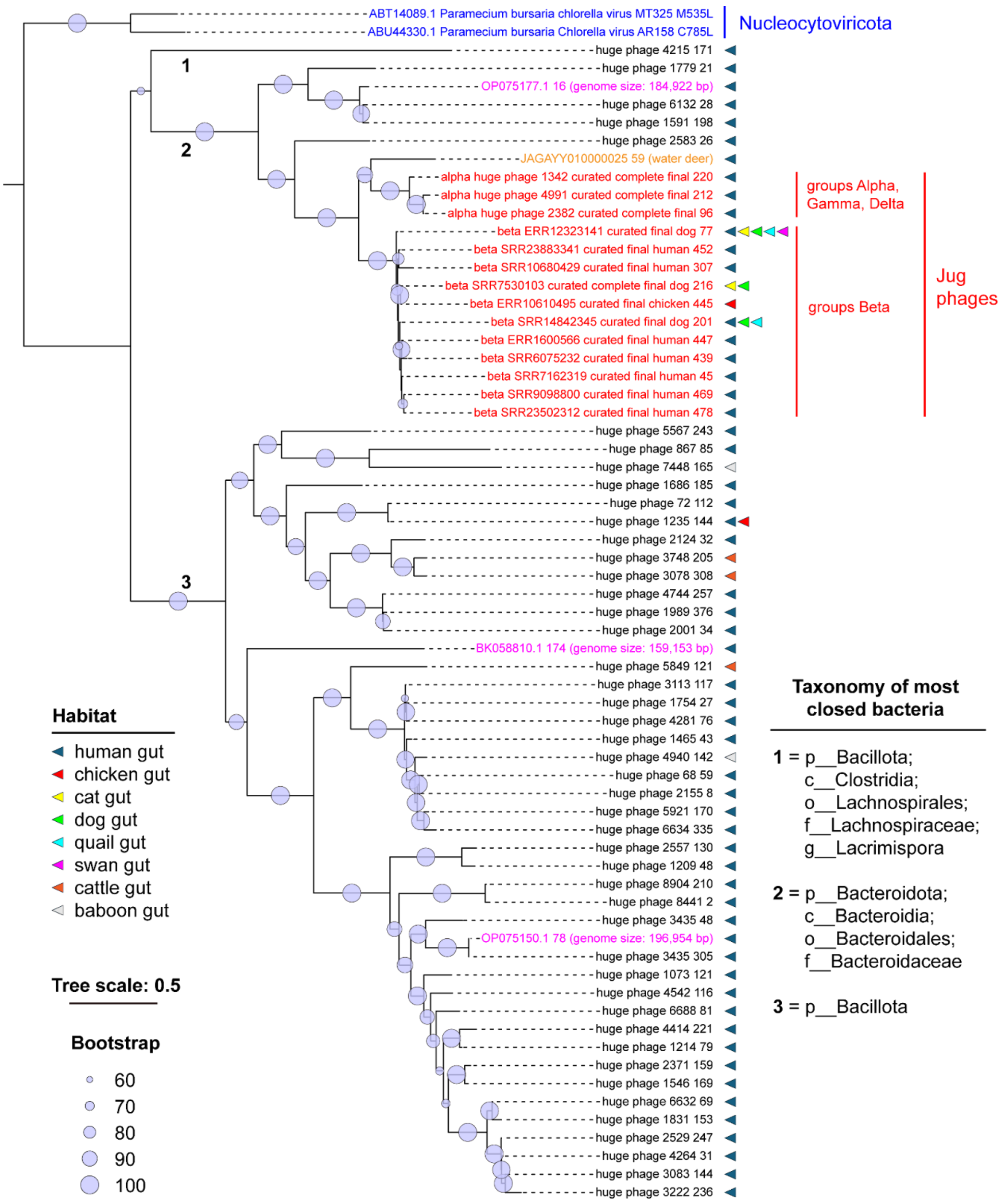


**Supplementary Figure 30 | The phylogeny of virus-encoded calcium-translocating p-type ATPases.** The ATPase protein sequences from the *Chlorella* viruses (M535L and C785L; in blue), the Jug phages (in red), the Jug phage relative from the water deer gut (in orange), the smaller-sized phages (NCBI Genbank; in pink), and other HPGC phages (in black) were included. Those sequences from Jug phages, NCBI Genbank, and other HPGC phages were respectively dereplicated using CD-HIT at 99% identity, and only the cluster representatives were included. The tree was rerooted using the two sequences from *Chlorella* viruses. A colored triangle indicated the animal host of the phages of each cluster. The most closed related taxonomy of the ATPases from smaller-sized phages (NCBI Genbank) and other HPGC phages was determined by searching them against the UniProt database.

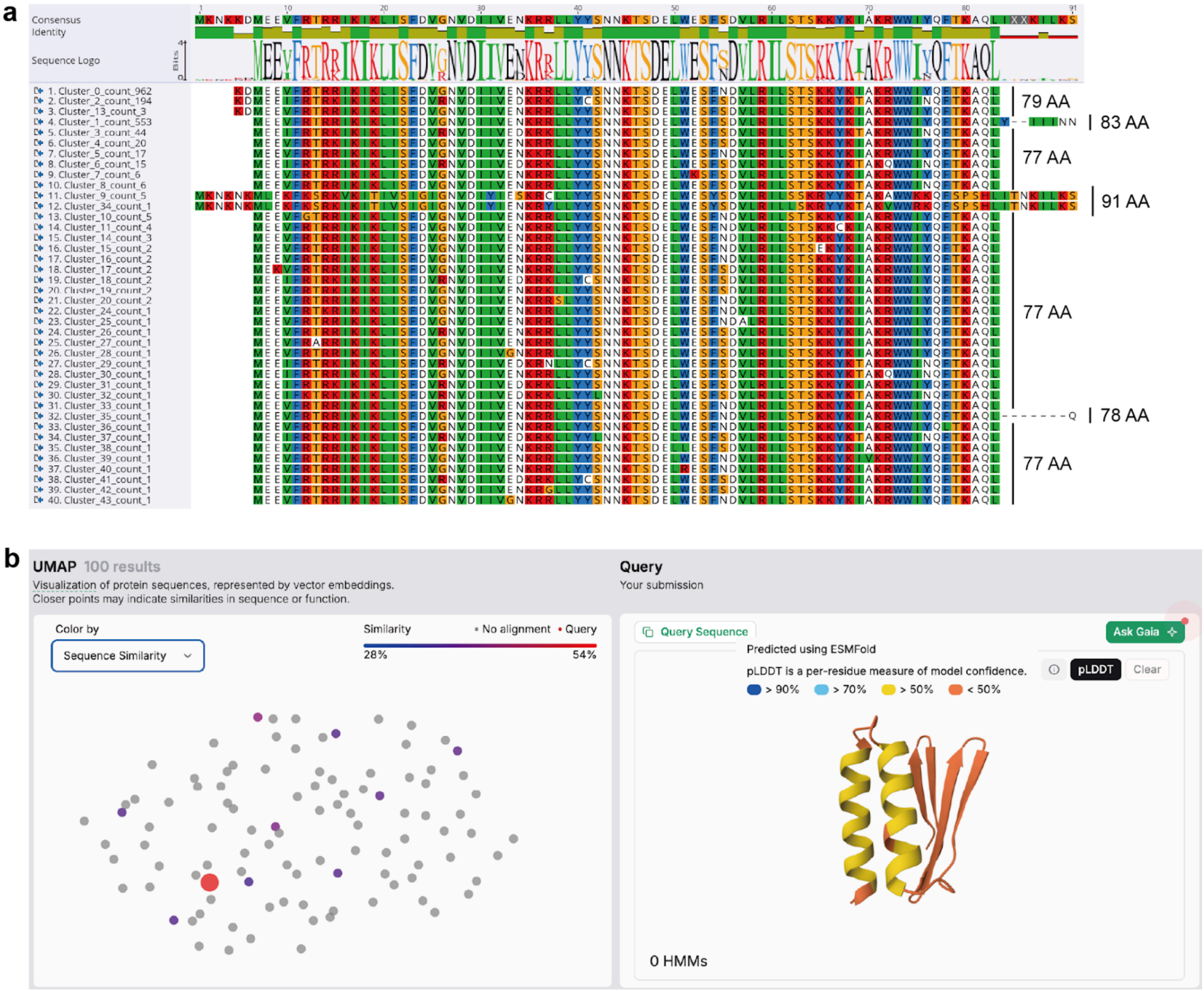


**Supplementary Figure 31 | The co-transcribed adjacent upstream gene of the calcium-translocating P-type ATPase gene encoded by the Jug phages.** (a) The protein sequence alignment of the adjacent upstream genes. The genes were from all curated and Logan-retrieved Jug phage genomes by BLASTp, and were dereplicated using CD-HIT (100% identity). Only the CD-HIT clusters are shown here (4 clusters were only partial genes and thus excluded). The cluster IDs and the total count of members in the cluster are shown on the left. The length of each representative is shown on the right. (b) The analyses of the adjacent upstream gene (the 77 AA version) were conducted using the recently published Gaia (Genomic AI Annotator) sequence annotation platform. The left panel shows the sequence similarity of the Ca$^{2+}$ ATPase adjacent gene (the big red circle) to those in the Gaia database; the right panel shows the predicted protein structure of the adjacent gene, with the pLDDT shown for each residue.
